# The dependence of shugoshin on Bub1-kinase activity is dispensable for the maintenance of spindle assembly checkpoint response in *Cryptococcus neoformans*

**DOI:** 10.1101/2024.02.28.582509

**Authors:** Satya Dev Polisetty, Krishna Bhat, Kuladeep Das, Ivan Clark, Kevin Hardwick, Kaustuv Sanyal

## Abstract

During chromosome segregation, the spindle assembly checkpoint (SAC) detects errors in kinetochore-microtubule attachments. Timely activation and maintenance of the SAC until defects are corrected is essential for genome stability. Here, we show that shugoshin (Sgo1), a conserved tension-sensing protein, ensures the maintenance of SAC signals in response to unattached kinetochores during mitosis in a basidiomycete budding yeast *Cryptococcus neoformans*. Sgo1 maintains optimum levels of Aurora B kinase Ipl1 and protein phosphatase 1 (PP1) at kinetochores. The absence of Sgo1 results in the loss of Aurora BIpl1 with a concomitant increase in PP1 levels at kinetochores. This leads to a premature reduction in the kinetochore-bound Bub1 levels and early termination of the SAC signals. Intriguingly, the kinase function of Bub1 is dispensable for shugoshin’s subcellular localization. Sgo1 is predominantly localized to spindle pole bodies (SPBs) and along the mitotic spindle with a minor pool at kinetochores. In the absence of proper kinetochore-microtubule attachments, Sgo1 reinforces the Aurora B kinaseIpl1-PP1 phosphatase balance, which is critical for prolonged maintenance of the SAC response.

## Introduction

The precise segregation of duplicated chromosomes in mitosis depends on the stable attachment of kinetochores with microtubules (MTs) and proper orientation on the mitotic spindle. The kinetochore is a multi-subunit protein complex composed of inner and outer layers (Musacchio and Desai, 2017). The inner kinetochore assembles onto centromeric chromatin and is constitutively associated with DNA. Subsequently, the outer kinetochore assembles and promotes the attachment of MTs emanating from microtubule organizing centers: centrosomes in animals or spindle pole bodies (SPBs) in yeast species. Apart from playing a role in MT attachment, the kinetochore also acts as a signaling platform for recruiting mitotic surveillance components that detect unattached or improperly attached kinetochores (Lara-Gonzalez et al., 2021, Zich and Hardwick, 2010, Musacchio and Salmon, 2007). Once errors are detected, cells remain arrested at metaphase until rectified.

Mitotic surveillance systems monitoring erroneous kinetochore-MT attachments include the spindle assembly checkpoint (SAC) and a chromosome passenger complex (CPC) component, Aurora BIpl1. The SAC comprises Mad1, Mad2, BubR1, Bub3, and the kinases Bub1 and Mps1 (Lara-Gonzalez et al., 2021). The SAC machinery conveys a ‘wait anaphase’ signal (MCC, the mitotic checkpoint complex) during cell division in response to unattached kinetochores that inhibits the anaphase-promoting complex (APC/CCdc20) from ubiquitylating B-type cyclins and securin (Hardwick et al., 2000, Musacchio, 2015, Sudakin et al., 2001). Upon the formation of proper kinetochore-MT attachments, the SAC is silenced. APC/CCdc20 inhibition is relieved, resulting in the degradation of B-type cyclins and securin, thus promoting mitotic exit and sister chromatid separation (Cohen-Fix et al., 1996, King et al., 1996).

Tension across sister-kinetochores attached to MTs is important in ensuring high fidelity of chromosome segregation. Tensionless kinetochore-MT attachments, if not rectified, result in the missegregation of chromosomes. Aurora BIpl1 acts as a tension sensor, detecting tension across sister kinetochores (Biggins and Murray, 2001, Tanaka et al., 2002, Hauf et al., 2003, Pinsky et al., 2006). In response to tensionless kinetochore-MT attachments, phosphorylation of outer kinetochore by Aurora BIpl1 results in the destabilization of kinetochore-MT interactions, leading to the generation of unattached kinetochores (Cheeseman et al., 2006, Ciferri et al., 2008) indirectly activating the SAC. Several reports implicate that Aurora BIpl1 can also directly activate the SAC machinery in response to unattached kinetochores, indicating that the function of Aurora BIpl1 is not just limited to detecting tensionless attachments (Kallio et al., 2002, Petersen and Hagan, 2003, Saurin et al., 2011, Santaguida et al., 2011, Pinsky et al., 2009). The kinase activity of Aurora BIpl1 also counteracts PP1 phosphatase recruitment to kinetochores (Liu et al., 2010). The activity of PP1 at kinetochores is required to terminate the SAC signaling and to help cells proceed into anaphase. PP1 dephosphorylates kinetochore substrates and thus promotes the stabilization of kinetochore-MT attachments (Lara-Gonzalez et al., 2021). Premature recruitment of PP1 to kinetochores results in early termination of SAC signals and leads to chromosome missegregation (Roy et al., 2019).

Shugoshin, another tension sensor, also detects tensionless kinetochore-MT attachments and promotes the biorientation of sister kinetochores during mitosis. Unlike Aurora BIpl1, shugoshin cannot directly destabilize tensionless attachments but acts as an adaptor protein in recruiting Aurora BIpl1, thus promoting the biorientation of sister kinetochores. Though shugoshin plays an important role in the biorientation of chromosomes, several reports indicate that during mitosis, it is dispensable for the SAC activation (Indjeian et al., 2005, Vanoosthuyse et al., 2007, Sane et al., 2021). Shugoshin primarily localizes to centromeres, and phosphorylation of histone H2A at T120 in humans/S121 in yeasts by the SAC component Bub1 kinase plays a central role in targeting it to centromeres (Kawashima et al., 2010, Fernius and Hardwick, 2007).

*Cryptococcus neoformans,* a budding yeast, diverged from other ascomycete yeasts such as *S. cerevisiae* and *S. pombe* at least 500 MYA (Heitman, 2011). *C. neoformans* is an opportunistic pathogen belonging to Basidiomycota and is considered a critical priority fungal pathogen by the World Health Organization (WHO). Several species of *Cryptococcus* cause fungal meningitis (Kronstad et al., 2011), a fatal infection in immunocompromised patients. *C. neoformans* undergoes semi-open mitosis, and its nuclear division occurs in the daughter bud (Kozubowski et al., 2013). This is strikingly different as compared to relatively well-studied yeasts such as *S. cerevisiae* and *C. albicans* where the nuclear division occurs at the mother-daughter bud-neck junction (Sutradhar et al., 2015). The kinetochore architecture is significantly rewired *in C. neoformans,* where a linker protein bridgin presumably replaced the loss of most inner kinetochore proteins (van Hooff et al., 2017, Sridhar et al., 2021). Moreover, metazoan-like kinetochore maturation dynamics in *C. neoformans* during cell cycle progression is distinctly different from most other budding yeast species (Kozubowski et al., 2013). The cell cycle-associated dynamics of the centromere-kinetochore complex in *C. neoformans* differ as well. Unlike other well-studied yeasts such as *S. cerevisiae*, *S. pombe, and C. albicans,* where kinetochores are constitutively clustered, the kinetochores in *C. neoformans* exhibit differential clustering dynamics, where they are unclustered during interphase and cluster during mitosis.

Despite high medical importance, mechanisms monitoring kinetochore-MT attachments have not been extensively studied in basidiomycete yeasts due to lack of molecular tools. Studies to explore the roles of SAC components Bub1 (Madbub), Mps1, and Mad1 in the checkpoint response using *C. neoformans* have just begun (Leontiou et al., 2022, Aktar et al., 2024). Moreover, *C. neoformans* exhibits aneuploidy-mediated fluconazole resistance (Sionov et al., 2010). Studying factors that regulate kinetochore-MT attachments in this organism with rewired kinetochore architecture and metazoan-like mitotic events might shed light on understanding aneuploidy-driven drug resistance.

Here, we probed the role of an evolutionarily conserved protein, shugoshin, in *C. neoformans* in monitoring the kinetochore-MT attachments during mitosis. We propose that shugoshin performs this function by maintaining higher Aurora BIpl1 and lower PP1 levels at kinetochores during MT depolymerizing conditions. Unlike ascomycete yeasts, shugoshin predominantly localizes to SPBs, along spindle MTs and a minor pool at kinetochores during mitosis. Apart from the atypical localization, we provide evidence that support that shugoshin targeting to its subcellular sites is not dependent on the kinase activity of Bub1.

## Results

### Shugoshin is required for efficient metaphase arrest in response to microtubule depolymerizing drugs

*SGO1* (FungiDB ORF ID: CNAG_05516) encodes the only form of shugoshin that is expressed in *C. neoformans.* The putative 645-aa-long protein coded by *SGO1* has an N-terminal coiled-coil domain and a C-terminal basic SGO motif (Fig 1A; Supplementary Fig 1A and B). Multiple sequence alignment of amino acid sequences revealed the presence of conserved residues required for PP2A binding (Supplementary Fig 1B) (Xu et al., 2009). The SGO motif in *C. neoformans* shugoshin also possesses a conserved lysine residue (K382) that is required for the binding of shugoshin to phosphorylated histone H2A (Supplementary Fig 1B) (Kawashima et al., 2010, Liu et al., 2013, Fernius and Hardwick, 2007). Previous studies reported that shugoshin plays a role in ensuring proper kinetochore-MT attachments during mitosis (see reviews (Gutierrez-Caballero et al., 2012, Marston, 2015)). Mutants affecting kinetochore-MT interactions and proteins participating in the error correction machinery and the SAC, are hypersensitive to MT depolymerizing drugs. To test if Sgo1 contributes to the fidelity of mitotic chromosome segregation, we created *sgo1* null mutants in *C. neoformans.* While Sgo1 was found to be dispensable for growth, *sgo1* null mutant cells displayed sensitivity to a MT-depolymerizing drug, thiabendazole (TBZ), similar to that of SAC mutant *mad2* (Fig 1B). These results suggest that *sgo1Δ* mutant cells in the basidiomycete yeast *C. neoformans* have phenotypes similar to those of ascomycete yeasts such as *Saccharomyces cerevisiae*, *Schizosaccharomyces pombe* and *Candida albicans* (Indjeian et al., 2005, Kitajima et al., 2004, Sane et al., 2021).

**Figure 1.**
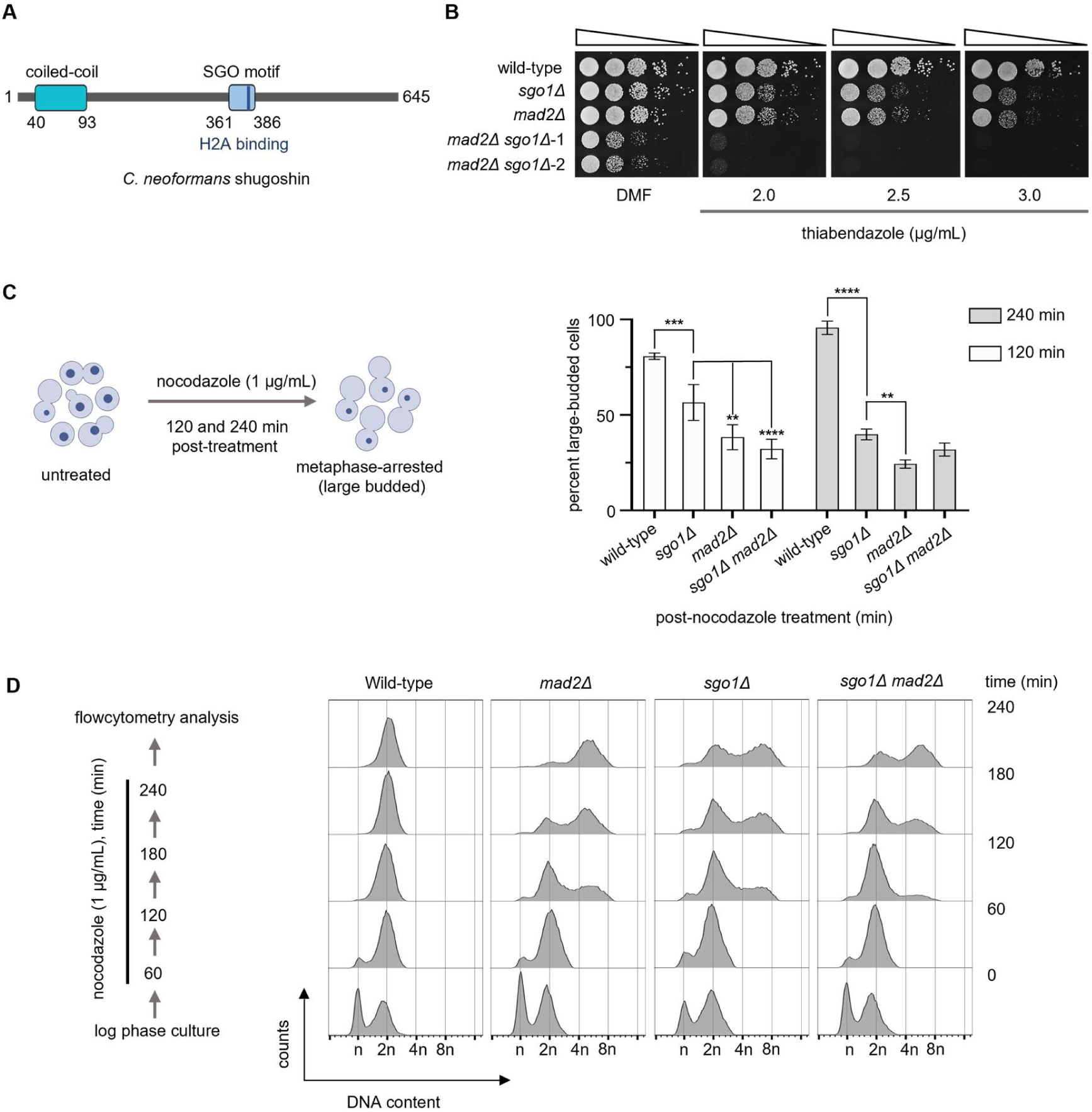
Shugoshin is required for a robust spindle assembly checkpoint arrest. **(A)** Schematic of the predicted motifs present in Sgo1 of *C. neoformans*. **(B)** A ten**-**fold serial dilution spotting assay to score for the sensitivity of strains CNVY108 (*SGO1 GFP-H4*), CNSD110 (*GFP-H4 sgo1Δ*), SHR741 (*GFP-H4 mad2Δ*), CNSD148 (*GFP-H4 sgo1Δ mad2Δ*) on plates containing the indicated concentrations of thiabendazole (TBZ). No drug represents DMF (dimethyl formamide) only. The plates were incubated at 30°C for 24 h. **(C)** *Left,* schematic of the assay to score for cell cycle arrest phenotypes when treated with nocodazole. Cells treated with nocodazole for 120 and 240 min were scored for cell cycle arrest phenotypes. *Right,* bar graphs representing the proportion of large-budded cells in the indicated strains, N=3, n>100, cells counted in each experiment. Two-way ANOVA with Tukey’s multiple comparison test was used to estimate the significance. ****p<0.0001, ***p<0.001, **p=0.0019/0.0072. Non-significant p-values were omitted from the graph. **(D)** *Left*, an outline of the assay to study the effect of nocodazole on the ploidy content of indicated strains using flow cytometry. *Right*, histograms depicting ploidy changes associated with nocodazole treatment.

Next, we sought to test if enhanced sensitivity to the MT poison observed in *sgo1Δ* mutant cells accentuate when combined with *mad2Δ* mutant. To test this, we created *sgo1Δ mad2Δ* double mutants. Such double mutant cells exhibited synthetic lethality in the presence of TBZ (Fig 1B), indicating that both Sgo1 and Mad2 are required for an efficient SAC response in MT-depolymerizing conditions. This striking synthetic lethality observed suggests that shugoshin is involved in functions such as biorientation apart from promoting the SAC response, which is lost in *mad2Δ*. If shugoshin is required for an efficient SAC response in *C. neoformans*, it is expected that the *sgo1Δ* mutant, when combined with a kinetochore mutant, would enhance sensitivities to TBZ. To explore this possibility, we used null mutant cells of a recently reported basidiomycete-specific kinetochore linker protein bridgin (Bgi1). It has been shown that *bgi1Δ* mutants are viable but exhibit gross chromosomal missegregation (Sridhar et al., 2021). *bgi1Δ* mutants, combined with *mad2Δ*, displayed enhanced sensitivity to TBZ compared to each of the single mutants alone (Sridhar et al., 2021). The spotting assay indicated that *sgo1Δ bgi1Δ* double mutant cells are hypersensitive to TBZ, similar to that of *mad2Δ bgi1Δ* (Supplementary Fig 2). Together, these results reveal that shugoshin plays a role in proper kinetochore-MT interactions and/or monitors defective attachments in response to spindle poison.

MT-depolymerizing drugs arrest wild-type cells at metaphase in the large-budded stage with an undivided nucleus. Only when the defects are rectified will such cells proceed to complete the cell division. To verify whether *sgo1Δ* mutants can induce metaphase arrest when MTs are depolymerized, we performed a microscopy-based assay to score for the cell cycle defects associated with *sgo1Δ* mutants by treating cells with nocodazole (Fig 1C). Nocodazole-treated wild-type cells of *C. neoformans* arrest at the large-budded stage with a visibly condensed chromatin mass (Fig 1C). After 240 min of nocodazole treatment, >90% of wild-type cells were at the large-budded stage with a condensed undivided nucleus (Fig 1C and Supplementary Fig 1C, D, and E). As expected, *mad2Δ,* a bonafide checkpoint-defective mutant, failed to arrest cells at the large-budded stage (Fig 1C). In the *sgo1Δ* mutants, though we initially observed an increase in the proportion of large-budded cells compared to *mad2Δ* mutants, after 240 min post-nocodazole treatment, the large-budded cell population significantly reduced (Fig 1C). Moreover, unlike wild-type, most large-budded cells in *sgo1Δ* mutants had an uncondensed chromatin mass similar to *mad2Δ* mutant cells (Supplementary Fig 1C, D, and E). In the case of *sgo1Δ mad2Δ* double mutants, no significant differences in the proportion of large-budded cells were observed when compared to *sgo1Δ* and *mad2Δ* single mutants (Fig 1C). Estimating the ploidy content in nocodazole-treated cells using flow cytometry (Fig 1D) revealed that wild-type cells arrested at 2n ploidy level. On the other hand, *mad2Δ* cells shifted towards 4n, indicating a failure in arresting cells at metaphase and initiating the next cell cycle events. Unlike *mad2Δ, sgo1Δ* cells displayed a ploidy content of 2n and 4n, revealing the presence of cells that are delayed at metaphase and cells that have initiated the next round of DNA replication. Similar results were also obtained in *sgo1Δ mad2Δ* double mutants. Taken together, we show that shugoshin is required for an efficient metaphase arrest in response to unattached kinetochores induced by MT-depolymerizing drugs.

### *sgo1Δ* mutant cells fail to maintain the SAC component Bub1 at kinetochores in response to prolonged microtubule depolymerization

Why are *sgo1Δ* mutant cells not arrested when treated with an MT poison at metaphase? It is possible that *sgo1Δ* mutant cells arrest at metaphase immediately after being treated with the MT poison but were unable to maintain the arrest upon prolonged exposure to the drug. To test this, we localized the SAC component Bub1 (Fig 2A) tagged with GFP in wild-type and *sgo1Δ* cells. A recent report (Leontiou et al., 2022) suggested that Bub1 localizes to kinetochores in *C. neoformans* and plays a central role in the SAC-mediated arrest. We synchronized *C. neoformans* cells by arresting them at G2 using hypoxia (Ohkusu et al., 2004) and released them in the presence of nocodazole (refer to Materials and Methods for further details) (Fig 2B). Inducing physical hypoxia in *C. neoformans* results in ‘unbudded’ G2 arrest. These G2-arrested cells enter mitosis synchronously when subjected to extensive aeration. Upon release into nocodazole-containing media, wild-type cells accumulated at the large-budded stage (Supplementary Fig 3) with GFP-Bub1 signals, which were maintained until the end of the experiment (Fig 2C and D). In the case of *sgo1Δ* cells, the proportion of Bub1-positive large-budded cells increased similarly to that of wild-type until 120 min. However, beyond 200 min post-nocodazole treatment, a sudden decline in the proportion of Bub1-positive large-budded cells was observed in the mutant (Fig 2C and D). Studies in *S. pombe* have shown that the kinase-dead version of Bub1 is checkpoint-proficient but fails to maintain prolonged SAC arrest (Fernius and Hardwick, 2007). A similar observation was reported in *C. neoformans*, where *bub1-kd* mutant cells failed to sustain the arrest induced by nocodazole (Leontiou et al., 2022). To understand how checkpoint responses differ in *sgo1Δ* from *bub1-kd,* we compared GFP-Bub1 localization dynamics in hypoxia-synchronized cells treated with nocodazole (Fig 2C and D). Though neither *sgo1Δ* nor *bub1-kd* mutant cells could maintain the arrest like the wild-type, *bub1-kd* cells were able to maintain the arrest longer than *sgo1Δ* cells.

**Figure 2.**
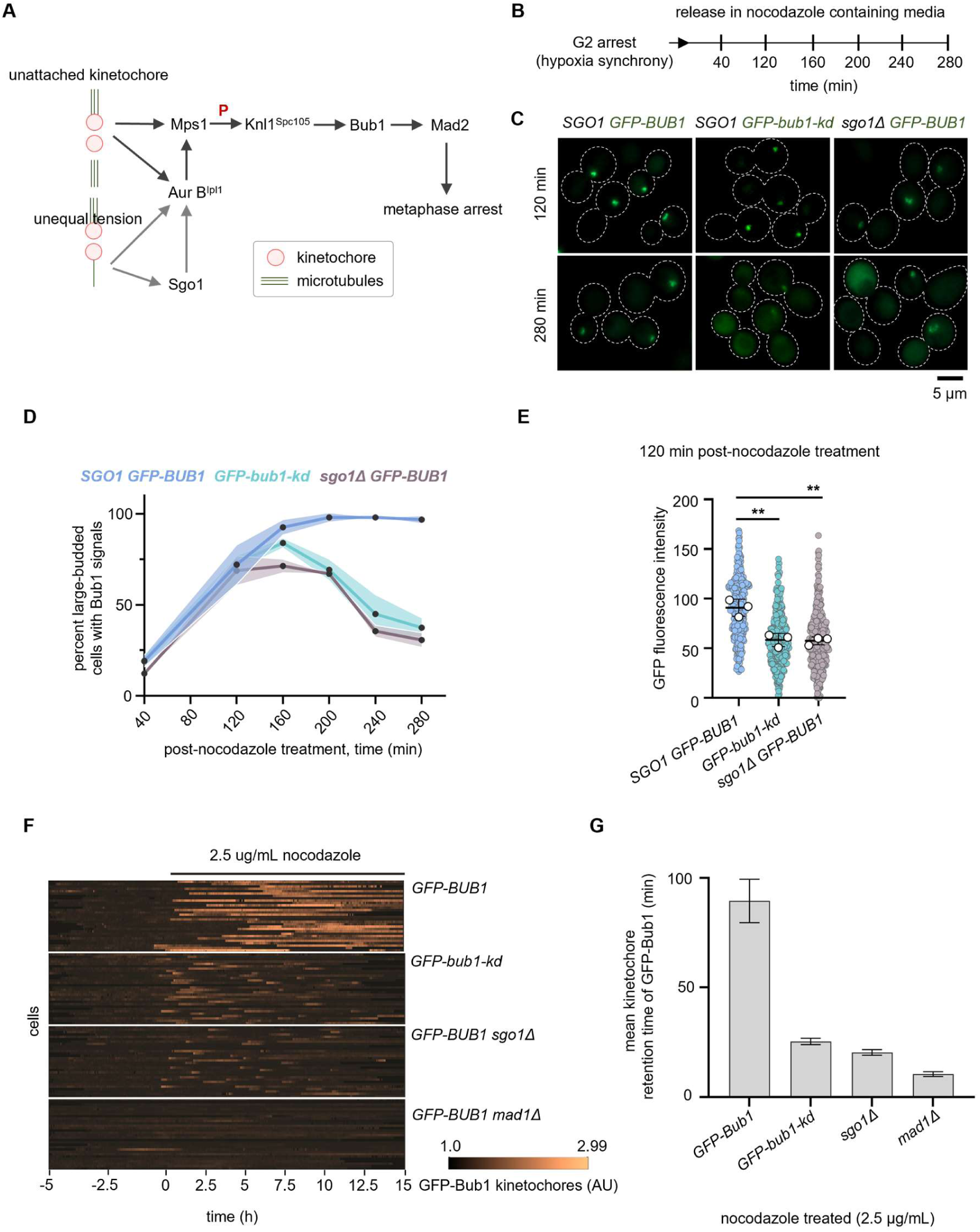
Shugoshin ensures Bub1 is retained at kinetochores to maintain the spindle assembly checkpoint arrest. **(A)** Schematic of the SAC pathway and the evolutionarily conserved role of Sgo1 and Aurora BIpl1 in detecting erroneous kinetochore-MT attachments. **(B)** An experimental workflow followed to determine the GFP-Bub1 localization dynamics under nocodazole-treated conditions. **(C)** Sub-cellular localization of GFP-Bub1 localization in IL102 (*SGO1*), IL143 (*bub1-kd*) and CNSD176 (*sgo1Δ*). For representation, cells treated with nocodazole for 120 and 280 min were shown. Scale bar, 5 μm. **(D)** Quantification of the proportion of large-budded cells with GFP-Bub1 signal in indicated strains treated with nocodazole. N=3, n>100 cells were analyzed, and the shaded regions above and below the line graphs represent error bands corresponding to SD. **(E)** Quantification of GFP-Bub1 fluorescence intensity values at 120 min post-nocodazole treatment. N=3, n>100 cells analyzed for each experiment. White circles represent the average intensity values of replicates. A mean of the average intensity values of replicates is represented as a black bar, and error bars represent SD. An unpaired t-test was used to test the significance, ** p=0.0092, ns =0.1131 **(F)** Microfluidics assays to determine the retention time of GFP-Bub1 at kinetochores in response to nocodazole treatment. Temporal heat maps of 30 randomly selected cells are shown. The heat map represents the changes in kinetochore localization signal of GFP-Bub1 over time. Each bright track on the *y*-axis of the heat map represents GFP-Bub1 signals from an individual cell (median of the brightest 5 pixels in each cell divided by the overall cell median brightness). The length of each bright track along the *x*-axis represents the time (min). The time of addition of nocodazole (2.5 μg/mL) is considered as 0 h. Images are taken every 2 min for 15 h. Assays were performed using strains IL102 (*GFP-BUB1*), IL143 (*GFP-bub1-kd*), CNSD176 (*GFP-BUB1 sgo1Δ*), and IL089 (*GFP-BUB1 mad1Δ*) strains. **(G)** Bar graphs representing the quantitative analysis of the GFP-Bub1 retention time at kinetochores obtained from microfluidics assays in the above-indicated strains. n=30, error bars represent SEM.

Moreover, Bub1 signal intensities in *sgo1Δ* and *bub1-kd* mutants were significantly reduced compared to the wild-type (Fig 2E). To further support these observations, we resorted to a single-cell microfluidics assay to quantitatively measure the checkpoint response in individual cells treated with the MT poison (Fig 2F and G). Using this system, we probed the GFP-Bub1 dynamics in wild-type, *sgo1Δ*, *bub1-kd, sgo1Δ bub1-kd,* and *mad1Δ* mutant cells treated with nocodazole. Similar to the results observed in the experiment performed with the synchronized cells, neither *sgo1Δ* nor *bub1-kd* mutant cells could maintain checkpoint arrest like wild-type as indicated by the lower mean kinetochore residence times of Bub1 (Fig 2G). Checkpoint defective *mad1Δ* mutant cells were used as a control. The *sgo1Δ bub1-kd* double mutant behaved like the *sgo1Δ* single mutant (Supplementary Fig 4A and B). These results reveal that *sgo1Δ* cells, similar to *bub1-kd* cells, are proficient in checkpoint activation but cannot maintain checkpoint signals during prolonged treatment with nocodazole.

### Aurora BIpl1 maintenance at kinetochores is affected in *sgo1Δ* mutants during prolonged checkpoint arrest

Shugoshin interacts with CPC proteins and helps in either the recruitment or maintenance of CPC components at centromeres/kinetochores in *S. cerevisiae*, *S. pombe* and humans (Hindriksen et al., 2017, Marston, 2015). Previously, we established the role of Aurora B^Ipl1^ (a CPC component) in regulating nuclear division during mitosis in *C. neoformans* (Varshney et al., 2019). Although it has been shown that the entry of Aurora BIpl1 into the nucleus coincides with the nuclear envelope breakdown (NEBD), its localization to kinetochores remained unexplored. In this study, we colocalized Aurora BIpl1 with the inner kinetochore protein CENP-A at different cell cycle stages (Supplementary Fig 5A). During G2, when kinetochores are clustered, we could only detect weak signals of Aurora BIpl1 that intensified as cells entered metaphase. Apart from the kinetochore localization, we could also observe spindle-like localization of Aurora BIpl1 during anaphase (Supplementary Fig 5A). Upon treatment with nocodazole, we observed colocalization of Aurora BIpl1 with kinetochores (Supplementary Fig 5B). To test whether kinetochore localization of Aurora BIpl1 is affected in *sgo1Δ* cells in *C. neoformans*, we synchronized both wild-type and *sgo1Δ* mutant cells by inducing physical hypoxia followed by a release in nocodazole-containing media (Fig 2B). We scored for Aurora BIpl1-3xGFP-positive large-budded cells. In *sgo1Δ,* we observed a reduced proportion of Aurora BIpl1- positive large-budded cells over time compared to wild-type (Fig 3A, B and Supplementary Fig 6). Similar to the Bub1 levels in *sgo1Δ* mutant cells, a reduction in the intensity of the Aurora BIpl1 signals in the *sgo1Δ* cells was also observed compared to the wild-type (Fig 3C). Taken together, we conclude that Sgo1 is required to maintain Bub1 and Aurora BIpl1 at kinetochores when MTs are depolymerized.

**Figure 3.**
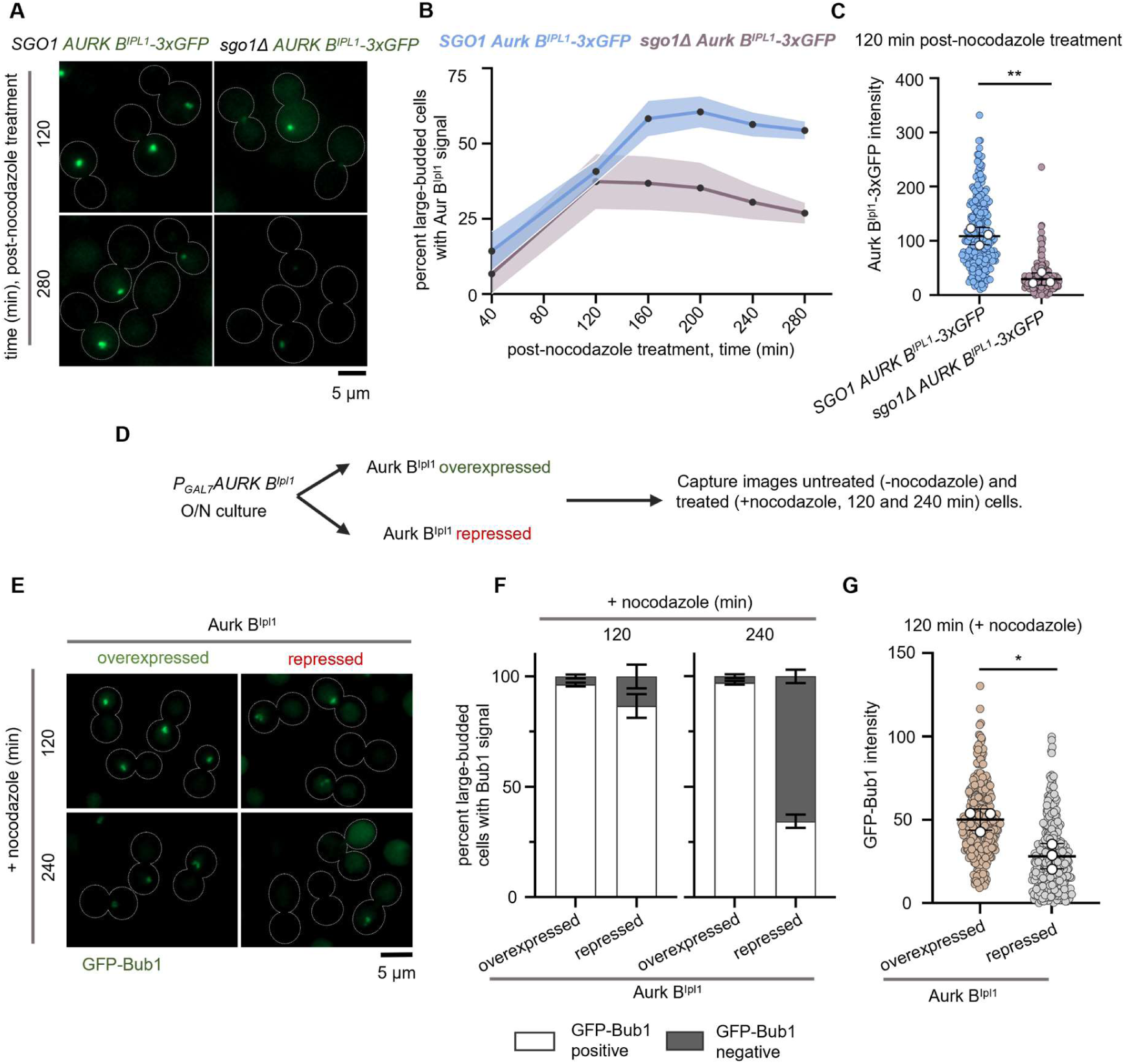
Shugoshin-dependent Aurora BIpl1 pool at kinetochores ensures maintenance of Bub1 to prolong the checkpoint arrest. **(A)** Sub-cellular localization of Aurora BIpl1-3xGFP localization in CNSD190 (*AURORA BIpl1-3xGFP mCherry-CENP-A SGO1*) and CNSD196 (*AURORA BIpl1-3xGFP mCherry-CENP-A sgo1Δ*) strains treated with nocodazole. For representation, cells treated with nocodazole for 120 and 280 min were shown. Scale bar, 5μm. **(B)** Quantification of the proportion of large-budded cells with Aurora BIpl1-3xGFP signals in the indicated strains. N=3, n>100 cells were analyzed, and the shaded regions above and below the line graph represent error bands corresponding to SD. **(C)** Quantification of Aurora BIpl1-3xGFP fluorescence intensity values at 120 min post-nocodazole treatment. N=3, n>58 cells for each experiment. White circles represent the average intensity values of replicates. The mean of the average intensity values of replicates is represented as a black bar. Error bars represent SD. An unpaired t-test was used to test the significance, ** p= 0.0013. **(D)** Schematic to assess the response of Aurora BIpl1 (over)expression or depletion to nocodazole treatment. **(E)** Sub-cellular localization of GFP-Bub1 localization in CNSD205 (*GAL7-3xFLAG-AURORA BIPL1 GFP-BUB1*) Aurora BIpl1 (over)expressed and repressed conditions treated with nocodazole. Cells treated with nocodazole for 120 and 240 min are shown for representation. Scale bar, 5 μm. **(F)** Bar graphs representing the proportion of large-budded cells with or without of GFP-Bub1 signals at 120 and 240 min of nocodazole treatment. N=3, n>100 cells analyzed for each experiment. Error bars represent SD. **(G)** Quantification of GFP-Bub1 fluorescence intensity values in Aurora BIpl1 (over)expressed and repressed conditions at 120 min of nocodazole treatment. White circles represent the average intensity values of replicates. The mean of the average intensity values of the replicates is represented as a black bar. Error bars represent SD. N=3, n> 79 cells analyzed for each experiment. An unpaired t-test was used to test the significance, *p=0.0439.

### Aurora BIpl1 is required for the maintenance of Bub1 at kinetochores in response to microtubule depolymerization

Reduced Aurora BIpl1 levels at kinetochores in *sgo1Δ* cells prompted us to test whether the absence of Aurora BIpl1 leads to failure of checkpoint arrest when treated with MT depolymerizing agents. For conditional depletion of Aurora BIpl1, we placed the gene under the *GAL7* promoter. We treated Aurora BIpl1 –(over)expressed (permissive, galactose media) and Aurora BIpl1-depleted (non-permissive, glucose media) cells with nocodazole (Fig 3D). We assayed for large-budded cells with a condensed chromatin mass as observed at metaphase. Cells (over)expressing Aurora BIpl1 and treated with nocodazole displayed a higher proportion of large bud arrested cells. In contrast, cells depleted of Aurora BIpl1 failed to arrest at the large-budded stage and exhibited an increase in the proportion of tri-budded cells (Supplementary Fig 7A and B). Moreover, approximately 50% of the large-budded cells displayed an uncondensed nuclear mass, similar to what was observed in *mad2Δ* mutants (Supplementary Fig 7A and B). These results suggest that Aurora BIpl1 is required for an efficient metaphase arrest when cells are treated with a spindle poison in *C. neoformans*.

To confirm these observations further, we examined GFP-Bub1 localization dynamics in cells depleted of Aurora BIpl1 and treated with nocodazole (Fig 3E). Aurora BIpl1- depleted cells subjected to nocodazole treatment for 120 and 240 min were able to load Bub1 initially but failed to maintain Bub1 at kinetochores upon prolonged treatment with nocodazole (Fig 3E and F), similar to what was observed in *sgo1Δ* mutant cells. We also noticed a significant reduction in the levels of Bub1 at kinetochores in Aurora BIpl1-depleted cells similar to that of *sgo1Δ* mutants (Fig 3G). These results reveal that Aurora BIpl1 in *C. neoformans* helps maintain Bub1 at kinetochores in the presence of nocodazole.

### Shugoshin antagonizes the recruitment of PP1 at kinetochores in response to nocodazole treatment

The kinase activity of Aurora BIpl1 and the phosphatase activity of PP1 are antagonistic at kinetochores (Tatchell et al., 2011, Lesage et al., 2011). We hypothesized that a reduction in Aurora BIpl1 levels leads to increased PP1 levels (Fig 4A), and higher PP1 levels may lead to the loss of Bub1 from kinetochores. We expressed PP1-GFP (FungiDB ORF ID CNAG_03706; Supplementary Fig 8) in a CENP-A-mCherry expressing strain (CNKB003). First, we studied the effect of *sgo1Δ* on PP1 localization during nocodazole treatment (Fig 4B). Both wild-type and *sgo1Δ* cells exhibited GFP-PP1 signals at kinetochores during the metaphase-arrested stage. Quantifying the fluorescence signal intensity of PP1 revealed a marginal but significant increase in the GFP-PP1 signal intensities at kinetochores in *sgo1Δ* compared to wild-type cells (Fig 4C). A similar result was observed when Aurora BIpl1-depleted cells were treated with nocodazole (Fig 4D and E). The SAC can be silenced by dephosphorylation of MELT repeats on Knl1Spc105 with the help of PP1. As a result, checkpoint components are removed from kinetochores (Pinsky et al., 2009, Rosenberg et al., 2011, Meadows et al., 2011, Espeut et al., 2012, Vanoosthuyse and Hardwick, 2009, Nijenhuis et al., 2014). Premature recruitment of PP1 to kinetochores is detrimental and leads to chromosome mis-segregation (Roy et al., 2019). Based on the results described above, we propose that the increase in PP1 levels at kinetochores leads to the loss of Bub1 signals in the absence of shugoshin or Aurora BIpl1, which interferes with the SAC maintenance in MT depolymerizing conditions.

**Figure 4.**
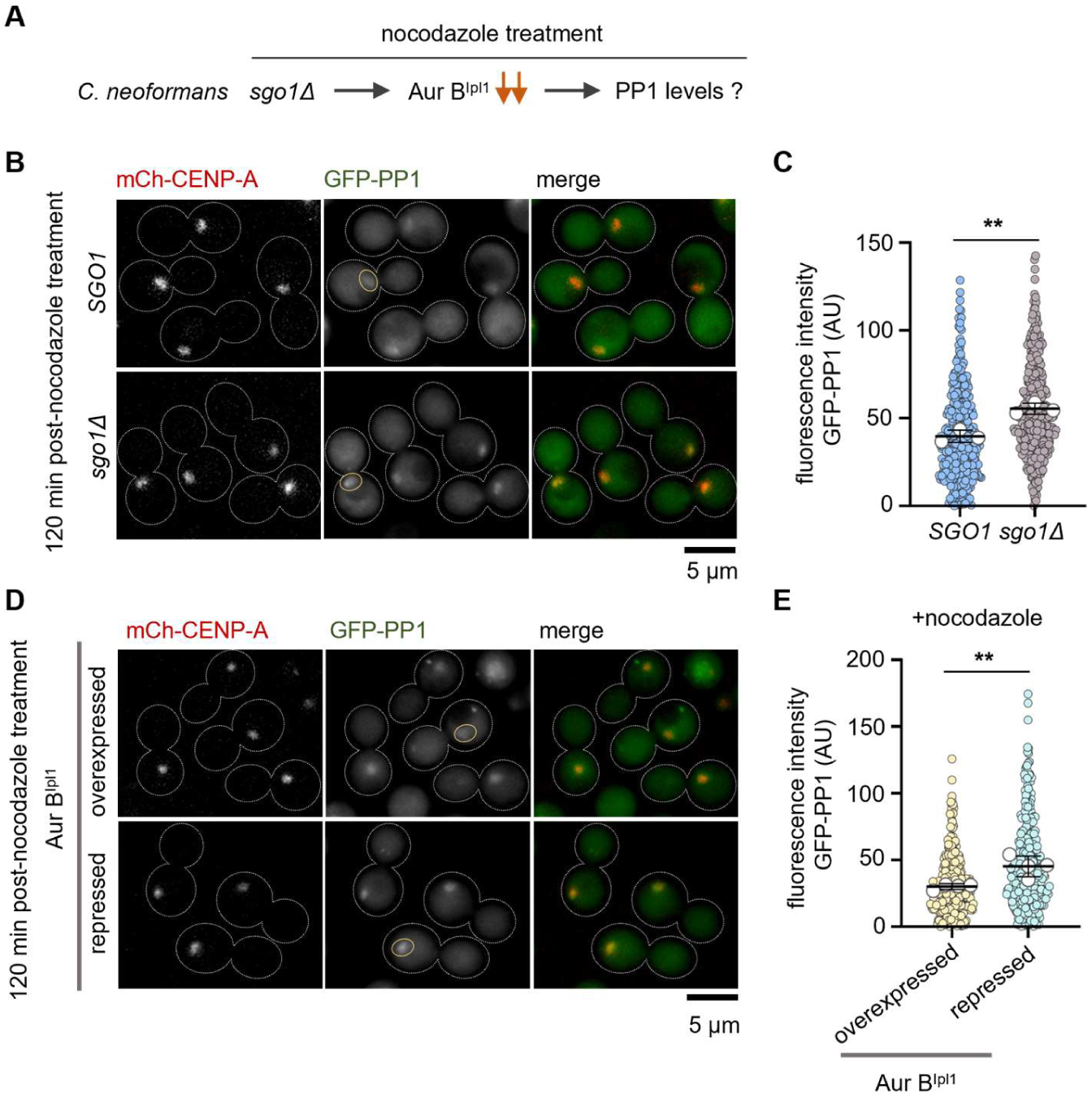
Sgo1-mediated Aurora BIpl1 recruitment prevents excess PP1 phosphatase at the kinetochore. **(A)** The effect of *sgo1Δ* on maintaining a balance between Aurora BIpl1 and PP1 levels at kinetochore in *C. neoformans*. **(B)** Sub-cellular localization of CNKB003 (*GFP-PP1 mCherry-CENP-A SGO1*) and CNSD180 (*GFP-PP1 mCherry-CENP-A sgo1Δ*) cells treated with nocodzole (1 μg/mL) for 120 min. Yellow ovals highlight PP1-GFP signals. **(C)** Quantification of GFP-PP1 fluorescence signal intensity values in the indicated strains treated with nocodazole. N=3, n>100 cells were analyzed for each experiment. White circles represent the average GFP-PP1 signal intensities calculated for each experimental repeat. An unpaired t-test was used to determine statistical significance. **p= 0.0044. The mean of the average intensity values of replicates is represented as a black bar. Error bars represent SD. **(D)** Microscopic images of CNKB023 (*GFP-PP1 mCherry-CENP-A GAL7-3xFLAG AURORA BIPL1*) Aurora BIpl1 (over)expressed and repressed conditions treated with nocodazole (1 μg/mL) for 120 min. **(E)** Quantification of fluorescence intensity values of GFP-PP1 signals in conditions mentioned above. N=4, n=100 cells for each experiment. White circles represent the average intensity values of replicates. Error bars represent SD. The mean of the average intensity values plotted is represented as a black bar. Paired t-test was used to determine statistical significance. **p= 0.0069.

### The tension-sensing function of shugoshin is conserved in *C. neoformans*

Shugoshin senses tensionless attachments by recruiting several factors that play a role in correcting tensionless attachments (Marston, 2015). Cohesin is a molecular glue that holds two sister chromatids together during mitosis until metaphase. Cohesin plays an essential role in generating tension that resists the pulling force of MTs, aiding high-fidelity chromosome segregation (Guacci et al., 1997, Michaelis et al., 1997). Loss of cohesion at centromeres results in tensionless kinetochore-MT attachments. To test if Sgo1 in *C. neoformans* detects tensionless attachments, we depleted one of the cohesin subunits, Scc1/Mcd1/Rad21, by placing it under the *GAL7* promoter both in the wild-type and *sgo1Δ* mutant cells. *C. neoformans* Scc1 (FungiDB ORF ID CNAG_01023) harbors N- and C-terminal Scc1 domains (Supplementary Fig 9A and B). We show that like *S. cerevisiae* and *S. pombe, SCC1* in *C. neoformans* is essential for viability (Supplementary Fig 9C). Upon depletion of Scc1, a high proportion of wild-type cells are arrested at the large-budded stage (Supplementary Fig 9D, E, and F). In contrast, the *sgo1Δ* mutants depleted of Scc1 did not show such an increase in the proportion of large-budded cells, indicating that shugoshin plays a role in detecting tensionless kinetochore-MT attachments and delays the cell cycle progression.

### Shugoshin is enriched at the spindle pole bodies (SPBs) and spindle during mitosis

Thus far, we reveal an unknown role of Sgo1 in maintaining Bub1 through the balance of Aurora BIpl1 and PP1 at kinetochores in *C. neoformans*. Next, we examined the cell cycle localization dynamics of Sgo1 in *C. neoformans*. Shugoshin localizes to the kinetochore during cell division in most species studied (Marston, 2015) (Supplementary Fig S11). To find the sub-cellular localization of Sgo1, we expressed GFP-Sgo1 from its native locus in the CENP-A-mCherry tagged strain. We synchronized the cells using hypoxia to examine the cell cycle stage-specific localization of Sgo1. No signals of Sgo1 could be observed during most of the interphase. In G2, where all centromeres cluster as a single punctum (Kozubowski et al., 2013, Sridhar et al., 2021), we observed Sgo1 signals near clustered centromeres (Fig 5A). Upon entry into mitosis, Sgo1 appeared as two distinct puncta that flanked the bar-like signals of CENP-A. Apart from the punctate signals of Sgo1, we observed weak signals along the pole-to-pole axis during metaphase and anaphase that resembled spindle-like localization. To examine whether Sgo1 localizes to spindle pole bodies (SPBs), we expressed Tub4-mCherry (γ-tubulin) in the GFP-Sgo1 expressing strain. GFP-Sgo1 signals colocalized with SPBs during G2 and mitotic stages (Fig 5B), indicating that Sgo1 localizes predominantly to SPBs during the mitosis in *C. neoformans*.

**Figure 5.**
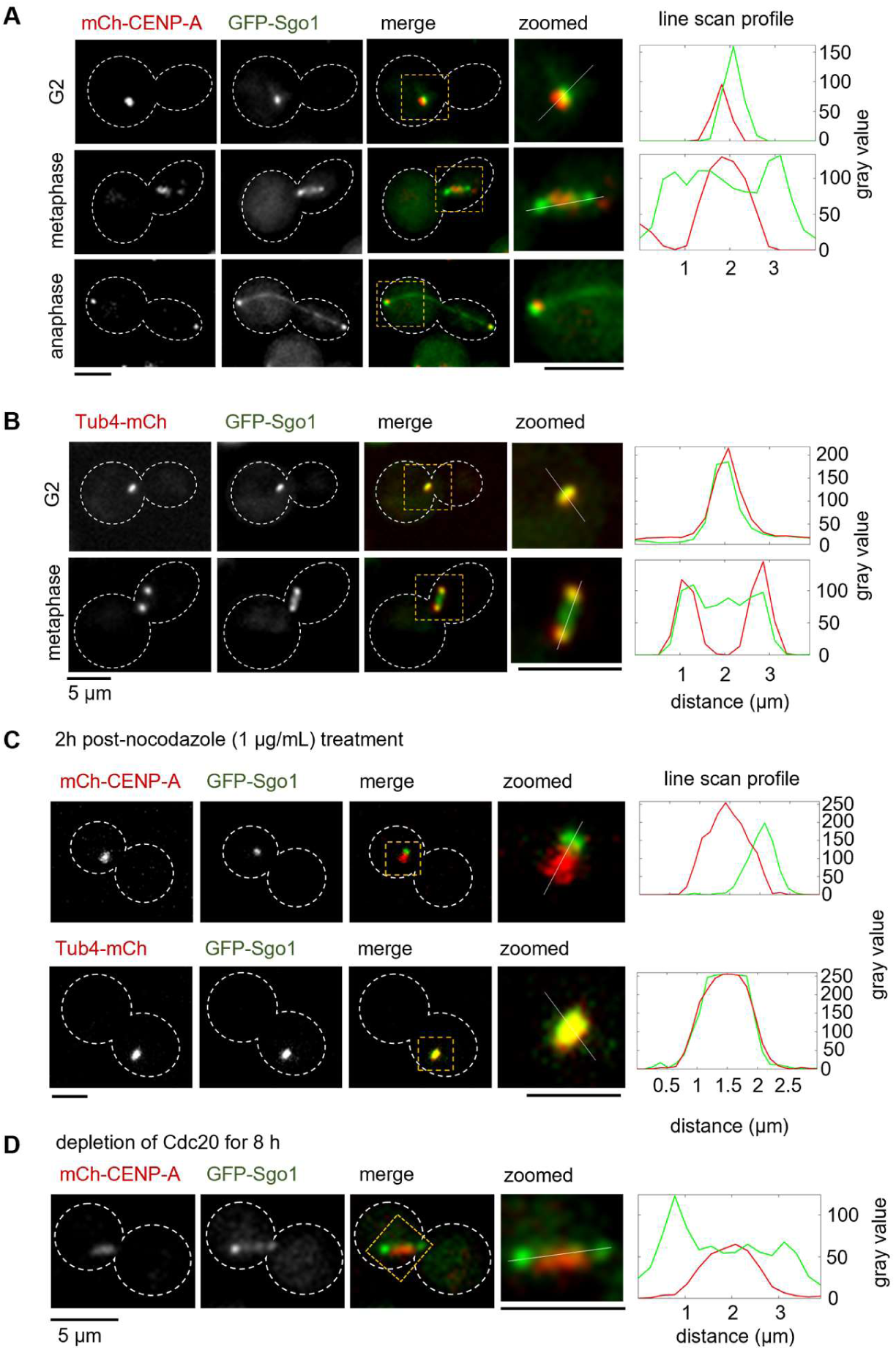
Kinetochore proximal association of Sgo1 in MT-depolymerizing conditions. (A-D) *Left,* microscopic images of GFP-Sgo1 localized with a kinetochore marker, CENP-A, and an SPB marker, Tub4, at different stages of the mitotic cell cycle, in nocodazole treated and Cdc20 depleted conditions. Scale bar, 5 μm. **(A)** CNSD125 (*GFP-SGO1 mCherry-CENP-A*). **(B)** CNSD130 (*GFP-SGO1 TUB4-mCherry*). **(C)** Nocodazole (1 μg/mL) treated condition. *Top panel*, CNSD125. *Bottom panel*, CNSD130. **(D)** Cdc20 depletion condition, CNSD167 (*GFP-SGO1 mCherry-CENP-A GAL7-3xFLAG-CDC20*). *Right*, line scan profiles of both GFP (Sgo1) and mCherry (CENP-A, Tub4) fluorescence signals. Distance plotted on the *x*-axis represents the length of the line drawn on the image to measure fluorescence intensity. Gray values on the *y*-axis represent the fluorescence signal intensities.

To test if the spindle-like localization during mitosis depends on the spindle MT integrity, we arrested cells at metaphase using two methods. First, we used nocodazole to depolymerize MTs that activate the SAC and arrest cells at metaphase. Second, we depleted Cdc20, an anaphase-promoting complex/cyclosome (APC/C) component. Depletion or inactivation of Cdc20 leads to metaphase arrest in *S. cerevisiae* (Hwang et al., 1998). Upon treating *C. neoformans* cells with nocodazole, we observed that Sgo1 colocalized with the SPB (Fig 5C). On the other hand, depletion of Cdc20 by placing it under the galactose regulatable promoter (*GAL7*) resulted in metaphase arrest, and the spindle-like localization of shugoshin along the pole-to-pole axis was maintained (Fig 5D). These observations led us to conclude that the spindle-like localization of Sgo1 requires intact MTs during mitosis.

### The kinase activity of Bub1 is dispensable for the function and localization of shugoshin during mitosis in *C. neoformans*

Studies from several yeast species and humans reveal that Bub1 is one of the major factors that recruit shugoshin to kinetochores. Although Bub1 plays a critical role, several other proteins such as cohesin, heterochromatin protein (HP1), CPC, and CENP-A also contribute to the targeting of shugoshin to kinetochores, see review (Marston, 2015) (Fig 6A). In the present study, to test if the Bub1 kinase activity required for shugoshin’s function is evolutionarily conserved in *C. neoformans*, we utilized the kinase-dead (*bub1-kd*) mutant of Bub1. *In vitro* assays reported earlier (Leontiou et al., 2022) revealed that Bub1 could phosphorylate the reconstituted human H2A containing nucleosomes, and kinase-dead mutants of Bub1 fail to phosphorylate. We performed a genetic interaction assay by scoring the TBZ sensitivity of *sgo1Δ*, *bub1-kd*, and *sgo1Δ bub1-kd* mutants (Fig 6B). If both Sgo1 and Bub1 kinase activity function in the same pathway in *C. neoformans*, one would expect little to no change in the sensitivity of double mutants compared to either of the single mutants, *sgo1Δ* and *bub1-kd*. By dilution spotting assay, we observed that the *bub1-kd sgo1Δ* double mutant cells behaved like that of the *sgo1Δ* mutant (Fig 6B). Moreover, the sensitivity of *sgo1Δ* mutant cells to TBZ is higher than that of *bub1-kd* mutant. There are two possibilities for the above observations. First, the enhanced sensitivity of the *sgo1Δ* mutant could be due to the additional functions of the shugoshin that do not rely on the kinase activity of Bub1. Second, the kinase activity of Bub1 does not play a significant role in dictating the kinetochore/kinetochore proximal localization of Sgo1. To test these possibilities, we have localized GFP-Sgo1 and performed a genetic interaction assay using mutants known to disrupt Bub1 kinase-mediated targeting of shugoshin in other species.

**Figure 6.**
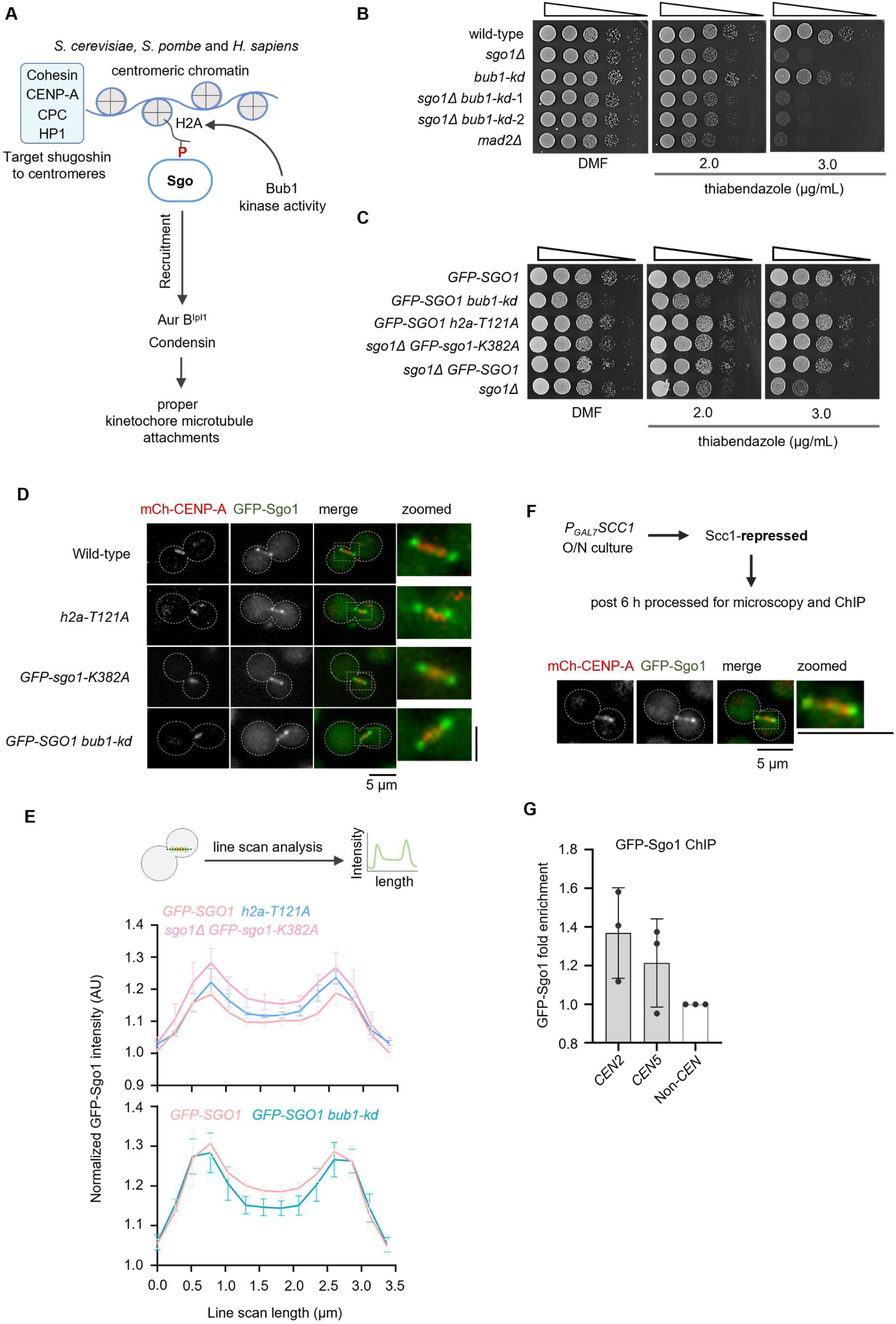
The kinase activity of Bub1 is dispensable for shugoshin-mediated maintenance of the spindle assembly checkpoint. **(A)** Schematic of the conserved pathway of shugoshin recruitment to kinetochores. **(B)** A ten-fold serial dilution spotting assay to score for the sensitivity of the CNSD110 (*sgo1Δ*), IL143 (*GFP-bub1-kd*), CNSD173 (*sgo1Δ GFP-bub1-kd*) double mutants and SHR866 (*mad2Δ*) to thiabendazole. **(C)** A ten-fold serial dilution spotting assay to score for the sensitivity of the CNSD125 (*GFP-SGO1 mCherry-CENP-A*), CNSD215 (*GFP-SGO1 mCherry-CENP-A bub1-kd-3XFLAG*), CNSD219 (*GFP-SGO1 mCherry-CENP-A h2a-T121A*), CNSD218 (*sgo1Δ GFP-sgo1-K382A-SH1 mCherry-CENP-A*), CNSD207 (*sgo1Δ GFP-SGO1-SH1*) and CNSD110 (*sgo1Δ*). **(D)** Microscopic images of GFP-Sgo1 localized with CENP-A at metaphase stage in wild-type and mutant backgrounds. Scale bar, 5 μm. (E) *Top,* Schematic showing the line scan analysis method adopted to score for the fluorescence intensity of GFP-Sgo1 along the spindle and SPBs. *Middle,* line scan profile of GFP-Sgo1 in the indicated strains synchronized using thiabendazole (10 µg/mL) treatment. *Bottom*, line scan profile of GFP-Sgo1 in the indicated strains synchronized using hypoxia. Distance on the *x*-axis represents the length of the line drawn on the image to measure fluorescence intensity. The *y*-axis represents the fluorescence signal intensity values. N=3, n>20 cells were analysed for each experiment. Each line graph represents mean fluorescence intensity values calculated from three experimental repeats and error bars represent SD. (F) *Top,* Schematic of the assay used to arrest cells at metaphase by depleting Scc1. *Below*, microscopic images of GFP-Sgo1 localized with CENP-A metaphase in Scc1 depleted conditions. **(G)** Fold enrichment of Sgo1 at kinetochores in metaphase arrested cell obtained by cross-linking chromatin immunoprecipitation (ChIP) followed by quantitative polymerase chain reaction (qPCR). *CEN2* and *CEN5* represent centromeres 2 and 5, non-CEN region represents *LEU2* locus. N=3, and error bars represent SD.

The lysine residue K382 in Sgo1 in *C. neoformans* corresponds to the lysine K492 in Sgo1 in humans (Supplementary Fig 1B). This residue is known to play a role in binding to phosphorylated H2A (Kawashima et al., 2010, Liu et al., 2013). A mutation in the lysine residue of shugoshin in fission yeast (K298) or humans (K492) results in the loss of kinetochore localization of shugoshin in mitosis (Liu et al., 2013, Kawashima et al., 2010). Similarly, mutating the S121 (budding/fission yeast) or T121 (humans) residue of histone H2A also abrogates the kinetochore localization of the shugoshin (Liu et al., 2013, Kawashima et al., 2010). To test the role of these conserved residues in *C. neoformans*, we mutated K382 in Sgo1 (Supplementary Fig 1B) and T121 in H2A to alanine (Supplementary Fig 10A). The plasmid constructs harboring either wild-type *SGO1* or a *sgo1*-*K382A* allele were integrated into the *SAFE HAVEN* locus (chromosome 1) (Arras et al., 2015) of the *sgo1Δ* strain. An overlap PCR construct harboring the *h2a-T121A* mutation was transformed into a strain expressing GFP-SGO1 and mCherry-CENP-A. If the kinase activity of Bub1 is indeed dispensable for shugoshin’s checkpoint function during mitosis, the strain carrying the *GFP-sgo1-K382A* allele would rescue the sensitivity to spindle poisons. Similarly, strain harboring the *h2a-T121A* allele would not show any sensitivity. Spotting assays on TBZ plates revealed that both sgo1*-K382A* and *h2a-T121A* mutants behaved like wild-type cells (Fig 6C). To further confirm these results, we localized Sgo1 in the above-mentioned mutant strains. Since Sgo1 exhibits both SPB and spindle-like localization during metaphase, we have obtained metaphase cells using hypoxia or thiabendazole arrest followed by release to score for the localization (refer to materials and methods for details). In both budding and fission yeasts, *bub1-kd*, *h2a-SA,* and *sgo-KA* mutants abolish kinetochore localization of shugoshin (Kawashima et al., 2010). In sharp contrast, we observed that *sgo1-K382A, h2a-T121A,* and *bub1-kd* mutants in *C. neoformans* did not abolish the SPB or spindle-like localization of Sgo1 during metaphase (Fig 6D and E). Sgo1 levels were unaffected in the above mutants (Supplementary Fig 10B). These results confirm that the kinase activity of Bub1 is dispensable for the recruitment of shugoshin and its SAC maintenance function in *C. neoformans*.

In *C. neoformans,* Sgo1 predominantly localizes to SPBs and the spindle during metaphase. The metaphase spindle localization of Sgo1 overlaps with CENP-A (Fig 5A). To probe if the kinetochores at the metaphase stage bind to Sgo1, we performed chromatin immunoprecipitation (ChIP) assay in Scc1-depleted cells (Fig 6F, top panel). Microscopy analysis of Scc1-depleted cells showed both SPB and spindle-like localization of Sgo1 (Fig 6F, bottom panel) as observed previously (Fig 5). ChIP assay indicates that there is only a minor enrichment of Sgo1 at centromeres in *C. neoformans* (Fig 6G), suggesting that the association of Sgo1 with centromeres could be transient in nature and/or only a minor pool of Sgo1 is associated with kinetochores and the majority could be along the spindle.

## Discussion

In this study, we characterized the mitotic functions of shugoshin in *C. neoformans*. The only form of shugoshin, coded by *SGO1*, predominantly localizes to SPBs and along the mitotic spindle in *C. neoformans*. In response to prolonged MT depolymerization, Sgo1 maintains adequate Aurora BIpl1 levels at kinetochores to antagonize premature phosphatase action of PP1 at kinetochores. Thus, by regulating an optimum balance of the protein kinase Aurora BIpl1 and the protein phosphatase PP1, shugoshin retains checkpoint responses until errors are corrected. Together, we report that Sgo1 regulates PP1 levels at kinetochores in response to microtubule depolymerization. We also demonstrated that the checkpoint maintenance function of Sgo1 is independent of the evolutionarily conserved Bub1 kinase activity-mediated targeting of Sgo1 to centromeres/kinetochores.

Sgo1 is not required to activate the SAC; rather, it helps prolong the SAC response by maintaining the levels of Bub1 at kinetochores in *C. neoformans*. Similarly, Aurora BIpl1 also helps in the maintenance of Bub1 levels at kinetochores in response to MT depolymerization conditions. Studies in *S. cerevisiae*, *S. pombe*, flies, and humans reveal that shugoshin interacts with the chromosome passenger complex (CPC) and is required for either recruitment or maintenance of Aurora BIpl1 to correct tensionless kinetochore attachments (Hindriksen et al., 2017, Marston, 2015). We show that apart from maintaining Bub1 levels at kinetochores, Sgo1 is also required to maintain Aurora BIpl1 levels when treated with nocodazole. Microfluidics and time course assays (Fig 2D, F, and 3B) indicate that *sgo1Δ* mutants can load Bub1 or Aurora BIpl1 at kinetochores albeit to a lesser level, but over time, the localization of these proteins is lost in the large-budded cells. This suggests that the loss of Bub1 or Aurora BIpl1 from kinetochores leads to failure in SAC maintenance in *sgo1Δ* cells during MT depolymerizing conditions. We hypothesize that Sgo1 regulates the SAC function through Aurora BIpl1 in two ways that may not necessarily be mutually exclusive: a) by maintaining high Aurora BIpl1 activity at kinetochores, thereby preventing the action of PP1 in terminating SAC signals, and b) by regulating Mps1 and Bub3/Bub1 (Saurin et al., 2011, Roy et al., 2022, Audett et al., 2022) at kinetochores, thereby modulating the SAC response. The fact that we observe an increase in the levels of PP1 at kinetochores in the reduced levels of Aurora BIpl1 indicates that Sgo1, more likely, acts through Aurora BIpl1 to antagonize the activity of PP1 at kinetochores thereby preventing the premature SAC termination.

The fact that Sgo1 is not required for the SAC activation but for the maintenance of the SAC response via maintaining a balance between Aurora BIpl1-PP1 at kinetochores also explains why *sgo1Δ mad2Δ* double mutants are synthetic lethal in the presence of MT depolymerizing drugs. The observed synergy indicates that both Sgo1 and Mad2 act in different pathways, where Sgo1 may play other roles such as biorientation of chromosomes that function (i) via the Aurora BIpl1-PP1-Bub1 axis; (ii) by recruiting condensin (Verzijlbergen et al., 2014, Peplowska et al., 2014) to the centromeres which biases sister kinetochores to bi-orient by affecting the kinetochore geometry; (iii) by maintaining the centromeric cohesion to prevent premature separation of sister chromatids, see review (Marston, 2015).

The mechanism of recruitment of shugoshin to kinetochores is a stratified process as it involves several proteins. Among these, Bub1 plays a critical role in targeting shugoshin to kinetochores in *S. cerevisiae*, *S. pombe*, *Xenopus laevis,* and humans (Liu et al., 2013, Kawashima et al., 2010, Boyarchuk et al., 2007) (Fig 6A). In humans, it has been reported that Cdk1-mediated phosphorylation of shugoshin promotes its interaction with cohesin and targeting to inner centromeres (Liu et al., 2013). Similarly, in *S. cerevisiae* (Kiburz et al., 2005) and *S. pombe* (Kitajima et al., 2004) cohesins are known to interact with shugoshin and this association in *S. cerevisiae* promotes the recruitment of shugoshin to pericentromeric regions in mitosis (Verzijlbergen et al., 2014, Kiburz et al., 2005). Apart from cohesins, it has been shown that a heterochromatin protein, HP1 promotes centromeric recruitment of shugoshin in *S. pombe* (Yamagishi et al., 2008). It has also been reported that shugoshin and the CPC complex mutually influence each other’s localization at centromeres. In *Drosophila*, the phosphorylation of MEI-S332 (shugoshin) by Aurora BIpl1 promotes its centromere localization (Nogueira et al., 2014, Resnick et al., 2006). Studies have also implicated the role of CENP-A and H3 nucleosomes in targeting shugoshin to centromeres as well (Wu et al., 2024, Mishra et al., 2018, Luo et al., 2016, Ng et al., 2013, Luo et al., 2010). Intriguingly, in C. neoformans, the checkpoint maintenance function of Sgo1 is not mediated through the Bub1-pH2A axis, as disruption of this pathway did not abolish the Sgo1 localization or the SAC maintenance function. This suggests that Bub1 is not a major player in recruiting CnSgo1 unlike the other species studied. We speculate that the Bub1 independent recruitment of CnSgo1 could be due to the non-canonical localization of the major pool of shugoshin at SPBs and along the mitotic spindle.

Altogether, this study illustrates the functional diversity of an evolutionarily conserved protein, shugoshin. A spatio-temporal balance of kinase and phosphatase activity is crucial for proper kinetochore-MT attachments and SAC signaling. Sgo1 monitors the kinetochore-MT attachment state in mitosis by balancing Aurora BIpl1 and PP1 levels at kinetochores. Based on the ChIP data, it is possible that there could be a transient and/or a minor pool of shugoshin associated with the kinetochores or kinetochore proximal regions. SPBs and spindle microtubules, along with other proteins such as HP1, CPC, CENP-A, and histone H3, might promote the kinetochore proximal association of Sgo1 in *C. neoformans*. This kinetochore proximal localization of shugoshin may not primarily rely on Bub1 kinase function. It is also possible that CPC or the activity of polo-like kinase (Cdc5/PLK1) as reported in humans (Wang et al., 2008) may also play a role in targeting Sgo1 to SPBs. Further investigation is needed to confirm the importance of SPBs/microtubules in regulating the function of Sgo1 in this organism of high medical importance.

## Materials and Methods

### Strains and media growth conditions

*C. neoformans* cultures were grown and maintained on YPD (1% yeast extract, 2% peptone, and 2% dextrose) and YPG (1% yeast extract, 2% peptone, and 2% galactose) media at 30°C unless specified. YPD+1M sorbitol or YPG+1M sorbitol plates were used for transformation. For the selection of transformants, 100 µg/mL of nourseothricin (NAT, clonNAT, Werner BioAgents), 180 µg/mL neomycin (G-418, Sigma-Aldrich), and 220 μg/mL of hygromycin (HiMedia) were used.

### Biolistic transformation

The biolistic transformation was done based on a previously described method (Davidson et al., 2000). Approximately 5 mL of *C. neoformans* culture was grown overnight, centrifuged at 4,000 rpm for 5 min, and resuspended in 300 µL sterile dH2O. Around 300 µL of cell suspension was spread on the YPD+1M sorbitol plate and dried. Gold micro-carrier beads (0.6 µm) coated with 2-5 µg of DNA were prepared, and 10 µL of 2.5 M CaCl2, 2 µL of 1M spermidine free base (Sigma-Aldrich) were added. The mixture was vortexed for 30 s, incubated for 10 min at room temperature, centrifuged, washed with 500 µL of 100% ethanol, and resuspended in 10 µL of 100% ethanol. The solution was placed on a micro-carrier disk and allowed to dry. Dried disks were placed in the biolistic transformation apparatus (Biolistic® PDS-1000/He Particle Delivery System). Helium gas pressure of approximately 1300 psi under a vacuum of a minimum of 25 inches of Hg was generated to bombard the cells with micro-carrier beads. The cells were incubated for 5-6 h in YPD+1M sorbitol medium at 30°C. Cells were then plated on YPD+NAT/NEO/HYG selection plate and incubated at 30°C for 3-4 days.

### Strains and plasmids

Lists of strains, plasmids, and primers used in this study are provided in Tables S1-S3. The construction of *C. neoformans* strains and plasmids is detailed below.

### GFP-SGO1 strains

The fluorescent fusion protein of shugoshin was generated by cloning the 1200 bp homology region and the *SGO1* promoter into the pVY7 plasmid containing the GFP and NAT sequences. GFP was tagged to the N-terminal region of the *SGO1* gene. The primers used for generating the clone are listed in Table S3. The 1200 bp homology region and a 500 bp *SGO1* promoter region were amplified from the *SGO1* gene using *C. neoformans* (H99) genomic DNA. The individual PCR fragments were gel purified. The amplified regions were cloned using BamHI, SacI, and NcoI sites. The final clone was linearized using HindIII and transformed into H99 and CNVY101. The transformants obtained were confirmed by PCR.

### *sgo1Δ* strains

An overlap PCR strategy was used to delete *SGO1* gene in the desired strains. Around 1000 bp region upstream to the start codon (ATG) and 1000 bp downstream region of the stop codon of the *SGO1* ORF was amplified using *C. neoformans* (H99) genomic DNA. The *NEO* resistance marker fragment was amplified from the pLK25 plasmid (Table S2). The individual PCR fragments were gel purified. Using these individual PCR amplified fragments, a final overlap cassette of ∼3800 bp was constructed using primers (Table S3). The final cassette was used to transform *C. neoformans*. The transformants obtained were confirmed by PCR. To delete the *SGO1* gene in different strain backgrounds, the *NEO* marker was swapped with either *NAT* or *HygB* genes amplified from the pVY7 and pSH7G plasmids (Table S2) using the same set of primers. The *sgo1Δ* cassette was used to transform H99, CNVY108, SHR741, SHR830, IL102, IL143, CNKB003 and CNSD190 strains. The transformants obtained were screened by PCR.

### TUB4-mCherry strain

An overlap PCR strategy was used to tag Tub4 with mCherry at C-terminus. Primers used for overlap cassette construction are listed in Table S3. Around 1000 bp region upstream to the stop codon (without including the stop codon) and 1000 bp 3’ UTR region after the stop codon was amplified from the *TUB4* ORF using *C. neoformans* (H99) genomic DNA. The mCherry sequence with antibiotic resistance marker NEO was amplified from the pLK25 plasmid (Table S2). The individual PCR fragments were gel purified. A final overlap cassette of ∼4500 bp was constructed using individual PCR fragments. The final cassette was transformed into CNSD127 strain. The transformants obtained were confirmed by PCR.

### GAL7-3xFLAG tagged SCC1, AURORA BIPL1, CDC20 strains

To place the desired gene under regulatable *GAL7p* along with an N-terminal 3xFLAG sequence, an overlap PCR strategy was used. Around 1000 bp regions upstream (US) and downstream (DS) of the start codon (ATG) of the ORFs of interest were amplified using *C. neoformans* (H99) genomic DNA. *GAL7* promoter with *HygB* resistance gene was amplified from pSH7G plasmid, and the 3xFLAG sequence was introduced using a primer (Table S3). The individual PCR fragments were gel purified. A final overlap cassette was assembled using the individual purified fragments. The cassettes generated were transformed into *C. neoformans* and the positive transformants were confirmed by PCR. The 3xFLAG tagged conditional mutants of *SCC1*, *AURORA BIPL1*, and *CDC20* generated using the strategy mentioned above are listed in Table S1.

### AURORA BIPL1-3xGFP mCherry-CENP-A strain

The H99 strain expressing Aurk BIpl1-3xGFP and mCherry-CENP-A was generated by the random integration of the plasmid pLKB71 (Table S2). The mCherry-CENP-A construct is expressed using the *H3* promoter. To create the strain, the plasmid backbone of pLKB71 was digested with HindIII and transformed into CNNV113, and the transformants were selected on the YPD-agar plate containing hygromycin selection. The colonies obtained were screened using a Zeiss Axio Observer 7 widefield microscope for the transformants positive for the mCherry-CENP-A signal.

### GFP-PP1 mCherry-CENP-A strain

The N-terminal GFP fusion protein of PP1 was generated by cloning around 1000 bp homology region and PP1 promoter into the pVY7 plasmid containing GFP and NAT sequences. Around 955 bp PP1 promoter region and 1000 bp PP1 homology region were amplified from PP1 ORF using *C. neoformans* (H99) genomic DNA. The PCR amplicons were gel purified. The digested amplicons of promoter and homology region were cloned into pVY7 using SacI, NcoI, and SpeI sites, respectively. The final clone was linearized using BglII and transformed into the CNVY101 strain, and the positive transformants were confirmed by PCR.

### GFP-sgo1-K382A-SH1, GFP-SGO1-SH1 *and* GFP-sgo1-K382A-SH1 mCherry-*CENP-A* strains

The N-terminal GFP fusion protein of wild-type and mutant (K382A) allele of *SGO1* was generated by cloning *SGO1* promoter, *SGO1* ORF, and 3’UTR into the pSD29 plasmid (Table S2) containing the GFP, HygB and *SAFE HAVEN* sequences. Around 2340 bp of *SGO1* ORF 315 bp of *SGO1* promoter, and 436 bp of 3’UTR were amplified from the *SGO1* gene using *C. neoformans* (H99) genomic DNA. The K382A mutation in the *SGO1* ORF was introduced using overlap PCR. The primers used are listed in Table S3. The individual PCR fragments were gel purified. The purified fragments of the *SGO1* promoter, *SGO1* ORF, and 3’UTR were cloned sequentially using SpeI, BamHI, HpaI, NheI, and ApaI sites. The final clone was linearized using XcmI and transformed into CNSD113. The transformants obtained were confirmed by PCR, and Sanger sequencing was performed to confirm the presence of the *K382A* mutation. The strain harbouring *GFP-sgo1-K382A* allele at *SAFE HAVEN* locus was transformed with mCherry-CENP-A-NAT plasmid linearized using BglII and the positive transformants were confirmed by microscopy.

### h2a-T121A GFP-SGO1 mCherry-CENP-A strain

A 1503 bp fragment of H2A ORF and 3’ UTR was amplified by PCR using *C. neoformans* (H99) genomic DNA and the T121A mutation was introduced into the 1503 bp fragment using overlap PCR strategy. A second fragment containing a Hygromycin marker was amplified from pSH7G. The homology region downstream of H2A 3’ UTR was amplified by PCR using *C. neoformans* (H99) genomic DNA. The primers used for generating the construct are listed in Table S3. The three PCR fragments were subsequently fused by overlap PCR. The final overlap product was used to transform the CNSD125 strain using the biolistic transformation method, and the transformants obtained were confirmed by PCR, and Sanger sequencing was performed to confirm the presence of mutation within the gene.

### bub1-kd 3xFLAG GFP-SGO1 mCherry-CENP-A *strain*

The 3xFLAG-tagged bub1-kd cassette was generated by cloning the 1200 bp homology region of BUB1 ORF harboring point mutations into the pSH7G plasmid containing the HYG sequences. The primers used for generating the clone are listed in Table S3. The 1200 bp homology region was amplified from the *bub1-kd* allele using genomic DNA obtained from the IL143 (*GFP-bub1-kd*) strain. The individual PCR fragments were gel purified. The amplified regions were cloned using SpeI and KpnI sites. The final clone was linearized using ApaI and transformed into CNSD125 background using the biolistic transformation method. The transformants obtained were confirmed by PCR.

### Thiabendazole sensitivity assay

For assaying sensitivity to thiabendazole, the O.D600 values of overnight cultures of wild-type and mutants used in this study were estimated, and serial dilutions starting from 105 cells to 101 cells were spotted on YPD-agar, DMF (Dimethyl Formamide); control, and YPD-agar plates with different concentrations of thiabendazole (2.0, 2.5, 3.0 and 4.0 μg/mL). The plates were incubated at 30°C for 24 to 48 h and imaged using a Bio-Rad chemidoc imaging system.

### Synchronization of *C. neoformans* by inducing physical hypoxia

For the synchronization of *C. neoformans* cells, we have used a protocol developed by Ohkusu et al., (Ohkusu et al., 2004) which utilizes a two-step protocol for the synchronization. *C. neoformans* cells, when grown in hypoxia conditions, result in unbudded G2 arrest, and cells release synchronously once shifted to extensive aeration conditions. Hypoxia synchrony is a two-step process. First, cells were grown at moderate aeration to 4.0 O.D600/mL in a 250 mL flask (Borosil Erlenmeyer flask with a screw cap was used for this experiment) with 50 mL of 1% YPD at 100 rpm, 30 °C. In the second step, the culture was diluted 1.5 times with fresh 1% YPD (final volume of 125 mL) and then incubated further for 5 h. The second step induces physical hypoxia, and at the end of the incubation, most cells get arrested at the unbudded G2 stage.

For the synchronization experiments, we used an overnight culture grown in 1% YPD to inoculate the secondary culture at 0.1 O.D600/mL in 1% YPD and grown to 1 O.D600/mL at 180 rpm, 30°C. This log phase culture was used for hypoxia synchrony, as mentioned above. The percentage of cells in the unbudded G2 stage was confirmed using a microscope. The culture was diluted to 1 O.D600/mL using fresh media, transferred to a sterile falcon or conical flask with appropriate culture volumes, and released under extensive aeration conditions (180 rpm, 30°C).

### GFP-Bub1 and Aurora BIpl1-3xGFP retention assay

To study the localization dynamics of GFP-Bub1 and Aur BIpl1-3xGFP in response to nocodazole treatment, wild-type, and *sgo1Δ* mutants were subjected to hypoxia synchrony, and the cells were released into 1% YPD media containing 1 μg/mL nocodazole. Time points were collected every 40 minutes until 280 min. The cells collected were imaged using a Zeiss Axio Observer 7 widefield microscope. A circular ROI of 12*12 or 6*6 pixels was used to measure the signal intensities. The presence or absence of GFP-Bub1 signals was scored manually. The cells that show bright punctate localization of Bub1 above the cell background were considered positive for Bub1.

### Imaging GFP-Sgo1 localization in *mCherry-CENP-A* and *TUB4-mCherry* strains

For imaging GFP-Sgo1 colocalization with CENP-A- and Tub4-mCherry tagged proteins, we synchronized cells using physical hypoxia. Cells were released into 1% YPD and collected every 10 min starting from 30 to 90 min. Cells were imaged using a Zeiss Axio Observer 7 widefield microscope.

### Cell cycle synchronization using thiabendazole (TBZ)

In order to obtain metaphase cells, the overnight-grown culture of *Cryptococcus* was inoculated at 0.2 O.D600/mL and grown to log phase. The log phase culture was treated with 10 µg/mL thiabendazole for 2 h. The cells arrested at G2/M were washed twice with 2% YPD and released into fresh pre-warmed 2% YPD by incubating at 30°C, 180 RPM for 2 min. The cells were harvested by spinning at 5000 RPM for 30 seconds, washed twice with sterile water, and imaged using a Zeiss Axio Observer 7 widefield microscope.

### Line scan analysis of GFP-Sgo1 signal at metaphase stage

For measuring the fluorescent intensity of Sgo1 at metaphase in wild-type and the *bub1-kd*, *h2a-T121A,* and *sgo1-K382A* strains, a line of 2-pixel thickness and a length of approximately 3.4 µm was drawn using a line tool available in Fiji (Schindelin et al., 2012). A single Z-plane containing both SPB and spindle signals was considered for intensity measurement. The fluorescence intensity values along the line were measured using the plot profile function. The intensity values of the GFP signal obtained from different cells were averaged and divided by the average background of the cells. The normalized intensity values were plotted using GraphPad Prism.

### Budding index calculation

The proportion of unbudded, small-budded, and large-budded cells was calculated based on the budding index (diameter of daughter cell/diameter of mother cell). The diameters of both mother and daughter cells were calculated along the mother-daughter axis using the line tool in Fiji (Schindelin et al., 2012).

### Fluorescence microscopy

For widefield microscopy, Zeiss Axio Observer 7 equipped with 100x Plan Apochromat 1.4 NA objective, Colibri 7 LED light source, motorized XYZ stage and PCO.edge 4.2 sCMOS camera was used. Images were acquired at 2x2 or 4x4 binning. Zen blue edition (2.3) software was used for controlling the microscope components and acquisition. For imaging, cells grown in culture media were washed twice with autoclaved dH2O and resuspended in water. The cell suspension was placed on microscope cover glass and imaged.

### Time-lapse microfluidic assays

The utility of microfluidics for single-cell analysis in *Cryptococcus* was demonstrated previously in studies of aging (Orner et al., 2019). We used the Alcatras cell traps (Crane et al., 2014) incorporated into devices allowing for use with multiple strains (Granados et al., 2018, Granados et al., 2017). Devices were moulded in polydimethylsiloxane (PDMS) from a SU8-patterned wafer with an increased thickness of 7 µm to accommodate the larger size of *C neoformans* cells compared to *S cerevisiae* (manufactured by Micro-resist, Berlin, design available on request). Imaging chambers for individual strains were isolated by arrays of PDMS pillars separated by 2 µm gaps. This prevents the intermixing of strains while cells experience identical media conditions. Before use, the devices were filled with synthetic complete (SC) media, supplemented with 0.2 g/L glucose, and containing 0.05% w/v bovine serum albumin (Sigma) to reduce cell-cell and cell-PDMS adhesion. Cells pre-grown to logarithmic phase in the same media (lacking the BSA) were injected into the device. An EZ flow system (Fluigent) delivered media at 10 µl per minute to the flow chambers and performed the switch to media containing nocodazole after 5 hours. This media also contained Cy5 dye to allow monitoring of the timing of the media switch. Image stacks were captured at 2<u>-minute</u> intervals at 4 stage positions for each strain, using a Nikon TiE epifluorescence microscope with a 60x oil-immersion objective (NA 1.4), a Prime95b sCMOS camera (Teledyne Photometrics) and OptoLED illumination (Cairn Research). Image stacks had 5 Z-sections, separated by 0.6 µm, captured using a piezo lens positioning motor (Pi).

### *Time-lapse* microfluidic assay image processing and analysis

Time-lapse assays were performed as described (Leontiou et al., 2022). Cell outlines were segmented using the baby algorithm (Pietsch et al., 2023). To quantify the kinetochore retention time of GFP-Bub1 fluorescence, we create a projection of the maximum values from all GFP sections, then divide the median fluorescence of the brightest 5 pixels within each cell by the median fluorescence of the cell as a whole. This ratio reliably measures protein aggregation at an organelle and has been extensively used for quantification (Cai et al., 2008, Hao and O’Shea, 2011, Lin et al., 2015). In the absence of nocodazole, cells with any outlier values for the ratio, defined as values greater than 3 scaled median absolute deviations away from the median, are removed. Cells present for fewer than 80% of recorded time points are also removed. The ratios were normalized for each strain by dividing the mean value for wild-type cells in the absence of nocodazole, measured in the same experiment. To calculate the kinetochore retention times, a threshold of 1.36 (Figure 2F and G) and 1.11 (Supplementary Fig 4A and B) was applied to the ratio values at each time point to determine the presence or absence of Bub1 at the kinetochore (Figure 2G and Supplementary Fig 4B). Small gaps in localization (due to kinetochore signal dropping briefly out of focus) were removed by applying a 1-dimensional morphological closing to the thresholded image, with a 3-pixel (3 timepoint or 6 minutes) structuring element. For the temporal heat maps (Figure 2F and Supplementary Fig 4A), the data for 30 cells present for the whole experiment are selected for inclusion from each strain pseudo-randomly using the Matlab rand function. There were a small number of outlier values within the heat maps, resulting from a fault in the LED control equipment. These were removed by replacing any values in the 5th percentile of a cell trace with the median value for all time points. The raw data and code used in data processing and plotting are available on request.

### Flow cytometry

5 mL of *C. neoformans* culture was grown overnight, and 2 O.D600 cells were taken and washed with sterile dH2O and fixed in 1 mL of 70% ethanol overnight at 4°C. Fixed cells were centrifuged at 4000 rpm and washed with 1 mL NS buffer(10 mM Tris-HCl pH 7.5, 250 mM sucrose, 1 mM EDTA (pH 8.0), 1 mM MgCl2, 0.1 mM CaCl2, and 7 mM β-mercaptoethanol) and resuspended in 200 µL of NS buffer added with 2 µg of RNase and incubated at 37°C for 3-4 h. Propidium Iodide (final concentration 12 µg/mL) was added to this mixture and incubated for 30 min at room temperature in the dark. 50 µL of the sample was suspended in 2 mL of 1x PBS (Phosphate buffered saline) solution, vortexed, and sonicated for 10 s at 30% Amp. 30,000 cells were analyzed by flow cytometry using FL2-A channel on (FACSAria III; BD Biosciences) at a rate of 500–2,000 events/s.

### Western Blotting

For immunoblotting, whole cell lysates were prepared by harvesting 10 O.D600 cells cultured using YPD. The cells were washed twice with sterile water and resuspended in 15% TCA (Trichloroacetic acid), and samples were frozen in a −20°C freezer. The cell suspension was thawed on ice and lysed using a mini-bead beater (Biospec product) with 1 min ON and 5 min OFF for three cycles at 3000 RPM. The cell lysate was centrifuged at 13000 RPM at 4°C for 10 min, and the supernatant was discarded. The pellet was resuspended in 2x sample buffer (0.1 M Tris HCl pH 6.8, 20% glycerol, 4% SDS, 715 mM β-mercaptoethanol and 0.2% bromophenol blue) and incubated for 5 min in a boiling water bath. The samples were cleared by spinning at 13000 RPM for 3 min, and the supernatant was loaded immediately onto an SDS-PAGE gel and separated. The proteins were transferred to a PVDF membrane (Bio-Rad) using the wet transfer method with a transfer time of two and a half hours at 90 V. The transfer efficiency was estimated by staining the blot with Ponceau S solution. The membrane was blocked using 3% skimmed milk and 2% BSA dissolved 1x PBST (0.1% Tween-20) solution at room temperature for 60 minutes. Post blocking, the blot was incubated in 2% skimmed milk and 1% BSA dissolved in 1x PBST solution containing anti-GFP at 1:3000 dilution overnight at 4°C. The membrane was washed thrice for 10 min with 1x PBS solution containing 0.1% tween-20. The blot was incubated with an anti-mouse secondary antibody (1:10000 dilution in 2% skimmed milk and 1% BSA dissolved in 1x PBST solution) for 1 hour at room temperature. The membrane was washed thrice for 10 min with 1x PBS + 0.1% tween-20, and the blot was rinsed in 1x PBS and was developed using Bio-Rad clarity ECL solution and imaged using Bio-Rad Chemidoc imaging system.

### Chromatin Immunoprecipitation (ChIP)

The ChIP protocol was adapted from (de Jonge et al., 2020, Mitra et al., 2014). ChIP was performed in metaphase-arrested cells by depleting Scc1. Overnight cultures of CNSD185 strain grown in 2% YPG (permissive media) were washed twice with 2% YPD and inoculated at 0.2 O.D600/mL in 150 mL of 2% YPD (Non-permissive) media. The cells were grown for 6 h at 30°C, 180 RPM. Post 6 h incubation, the extent of metaphase arrest was assessed using microscopy, and the culture was fixed by adding freshly prepared methanol-free formaldehyde at 3% final concentration and incubated at 25°C, 100 RPM for 35 minutes. The fixed cells were quenched with 1.5 M Tris HCl pH 8.0 for 5 mins at 25°C, 100 RPM. The cells were pelleted and washed with ice-cold water flash frozen using liquid Nitrogen, and stored at −80°C. The cells were lysed in a buffer containing 50 mM HEPES pH 7.4, 1% Triton-X, 140 mM NaCl, 0.1% Na-deoxycholate, 1 mM EDTA, 1x Protease Inhibitor Cocktail (Sigma) with 10 cycles of bead beating using a mini-beadbeater (BioSpec product) at 3500 RPM for 2 mins ON and 5 mins OFF on ice.

The lysate was transferred to a chilled 15 mL falcon on ice, and the cross-linked chromatin was fragmented using an ultrasonicator (Diagenode Bioruptor Pico) at high frequency, 30 s ON, 30 s OFF mode for 50 cycles to achieve the desired 250-500 size range. The sample was transferred to a chilled 2 mL tube, and centrifuged at 10000 RPM for 10 min at 4°C. The debris-free supernatant was carefully transferred to a fresh 2 mL tube and incubated with 20 µL of control trap beads (Chromo Tek) at 11 RPM for 3 h at 4°C. After incubation, bead-free supernatant was recovered by centrifugation at 500 RCF for 2 min, and GFP-trap and block beads were added to the respective tubes. The tubes were incubated at 11 RPM rotation for 12 h at 4°C. After incubation, the beads were recovered by centrifugation at 1000 RCF for 1 min at RT, and extensive washes were performed at RT with intermittent agitation at 11 RPM for 5 min. First, the beads were washed with 1 mL lysis buffer (50 mM HEPES pH 7.5, 140 mM NaCl, 1% Triton-X 100, 0.1% sodium deoxycholate, 2 mM PMSF), recovered by centrifugation at 1000 RCF for 1 min at RT and then washed 2 times each with the following buffers, with recovery at 1000 RCF for 1 min following each wash: (i) 20 mM Tris HCl pH 8.0, 200 mM NaCl, 2 mM EDTA, 1% Triton-X-100, 0.1% SDS. (ii) 20 mM Tris HCl pH 8.0, 500 mM NaCl, 2 mM EDTA, 1% Triton-X-100, 0.1% SDS. (iii) 20 mM Tris HCl pH 8.0, 250 mM LiCl, 2 mM EDTA, 1% NP-40, 1% sodium-deoxycholate, pH 8.0. (iv) TE: 10 mM Tris HCl pH 8.0, 1 mM EDTA. The IP complex was recovered by incubating the beads with 250 µL of elution buffer (10 mM Tris HCl pH 8.0, 1 mM EDTA, 1% SDS, 200 mM NaCl) at 65°C for 15 min with intermittent flicking. The process was repeated with another 250 µL buffer and pooled to obtain a 500 µL IP sample. The input volume was adjusted to 500 µL with an elution buffer. For de-crosslinking, 20 µL of 5 N NaCl was added to each sample and incubated overnight at 65°C. Next day, 20 µL 1M Tris-HCl pH 6.8, and 10 µL 0.5 M EDTA pH 8.0 were added to each tube and treated with Proteinase K (2 µL/sample) at 42°C for 1 h followed by RNase treatment (5 µL of 10 mg/mL RNase A/sample) at 37°C for 1 h. The DNA was extracted with an equal volume of Tris pH 8.0-saturated phenol:chloroform:isoamyl alcohol (25:24:1) and precipitated (with 1 mL of 100% ethanol with 100 µL 3 M sodium acetate pH 5.3) overnight at −80°C. The DNA was pelleted by centrifugation at 14000 RPM for 30 min at 4°C, washed with chilled 70% ethanol, air-dried, and finally resuspended in 20 µL TE pH 8.0. The ChIP-qPCR reactions were carried out in a total 10 µL volume using CEN and Non-CEN primers listed in Table S3.

### Statistics and microscope image analysis

P-values were assessed by unpaired, two-tailed t-test or two-way ANOVA with Tukey’s multiple comparison test using GraphPad Prism 8.00 (GraphPad software). Error bars represent the standard deviation (SD) or standard error of the mean (SEM) as mentioned for each experiment. N for each experiment is mentioned in the figure legends. Fiji software (Schindelin et al., 2012) was used to process and analyze microscope images. Fluorescence intensity values of signals were quantified by drawing a circular region of interest (ROI) of defined pixel size on the sum intensity Z-projected images (Figure 2E, 3C and G) or on single Z-plane images (Figure 4C and E). The intensity values were calculated by subtracting the product of mean background fluorescence intensity and area of the ROI from integrated density, and the resultant values were plotted using GraphPad Prism.

## Acknowledgments

We thank members of the Sanyal laboratory for their support, discussion, and suggestions on the work. We thank A. A. Jeyaprakash, Institute of Cell Biology, School of Biological Sciences, University of Edinburgh, UK, for the discussion and suggestions on the work. We thank I. Leontiou and T. Davies (University of Edinburgh) for *bub1-kd* strain construction. We thank the in-house flow cytometry facility at JNCASR and N. Nala for assistance with flow cytometric analysis. We thank V. Yadav, Duke University, USA, for the reagents provided for Tub4-mCherry strain construction. S.D.P. is a Research Associate supported by SERB funding by the Government of India. KB is a senior research fellow (SRF) supported by CSIR, Government of India. KD is senior research fellow (SRF) supported by JNCASR. KH was supported by the Leverhulme Trust (RPG-2018-379); IC was supported by the Wellcome Trust-University of Edinburgh Institutional Strategic Support Fund, and we thank P. Swain (University of Edinburgh) for leading on this ISSF funding. KS is a JC Bose National Fellow (JCB/2020/000021) of the Science and Engineering Research Board (SERB), Department of Science and Technology, Government of India. Kaustuv Sanyal acknowledges financial support from the Department of Biotechnology, Government of India (IC-12025(22)/3/2023-ICD-DBT), SERB (CRG/2023/001077), and intramural funding from JNCASR.

## Supplementary Figure legends

**Supplementary Figure 1.**
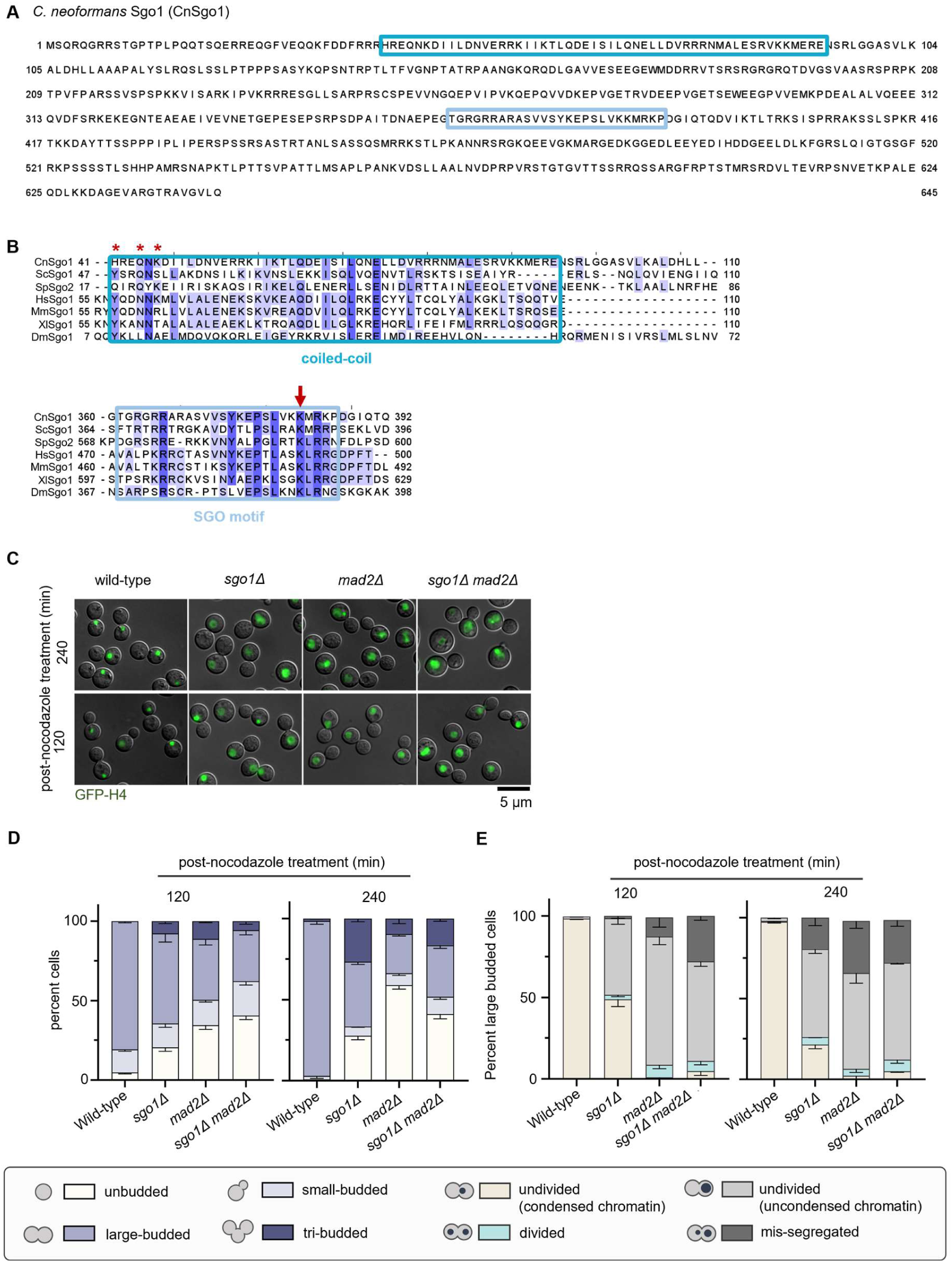
*C. neoformans* Sgo1, carrying a conserved coiled-coil domain and the SGO motif, is required to maintain SAC arrest in nocodazole-treated cells. **(A)** The protein sequence of *C. neoformans* Sgo1. **(B)** Multiple sequence alignment (MSA) of the CnSgo1 N-terminal coiled-coil and the C-terminal SGO motif with vertebrate and invertebrate species (Cn-*C. neoformans*; Sc- *Saccharomyces cerevisiae*; Sp- *Schizosaccharomyces pombe*; Hs- *Homo sapiens*; Mm- *Mus musculus*; Xl- *Xenopus laevis*; Dm- *Drosophila melanogaster*). Multiple sequence alignment of Sgo1 homologs was performed using Clustal Omega (Sievers et al., 2011), and the alignment was formatted using Jalview 2 (Waterhouse et al., 2009). *Represent amino acid residues involved in PP2A binding. The dark blue circle represents the key residue (K382 in *C. neoformans*) required for Bub1-mediated kinetochore proximal centromere localization of shugoshin. The teal blue colored box represents the coiled-coil domain. The light blue colored box represents the conserved basic SGO motif. **(C)** Microscopic images of GFP-H4 in CNVY108 (*SGO1 MAD2*), CNSD117 (*sgo1Δ*), SHR866 (*mad2Δ*), CNSD148 (*sgo1Δ mad2Δ*) treated with nocodazole (1 μg/mL) for 120- and 240-min. Scale bar, 5 μm. **(D)** Bar graphs representing the proportion of unbudded, small-budded, large-budded, and tri-budded cells. N=3, n>100 cells counted in each experiment. **(E)** Bar graphs representing the proportion of large-budded cells of indicated phenotypes, N=3, n>100 cells counted in each experiment.

**Supplementary Figure 2.**
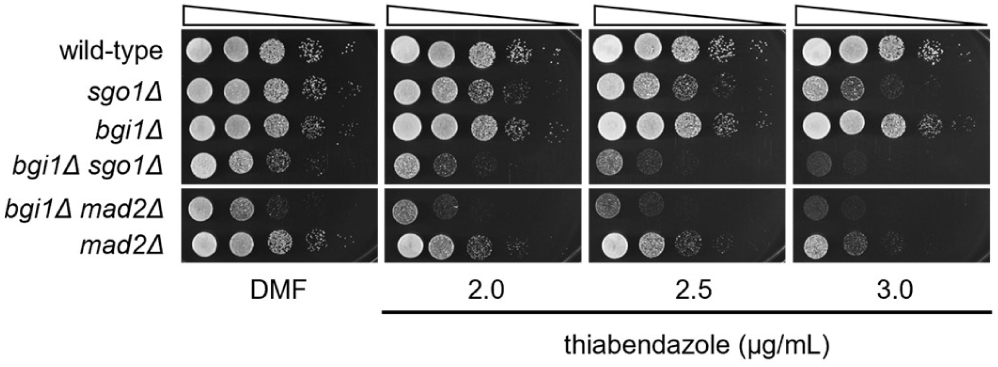
*sgo1Δ bgi1Δ* double mutant cells exhibit enhanced sensitivity to MT depolymerizing drug. A ten**-**fold serial dilution spotting assay to score for the sensitivity of the wild-type CNVY108 (*SGO1 BGI1 GFP-H4*), CNSD117 (*sgo1Δ GFP-H4)*, SHR741 (*mad2Δ GFP-H4*), SHR830 (*bgi1Δ GFP-H4*), CNSD148 (*sgo1Δ mad2Δ GFP-H4*) and CNSD163 (*sgo1Δ bgi1Δ GFP-H4*) to thiabendazole. No drug represents DMF (Dimethyl formamide) only. The plates were incubated at 30°C for 24 h.

**Supplementary Figure 3.**
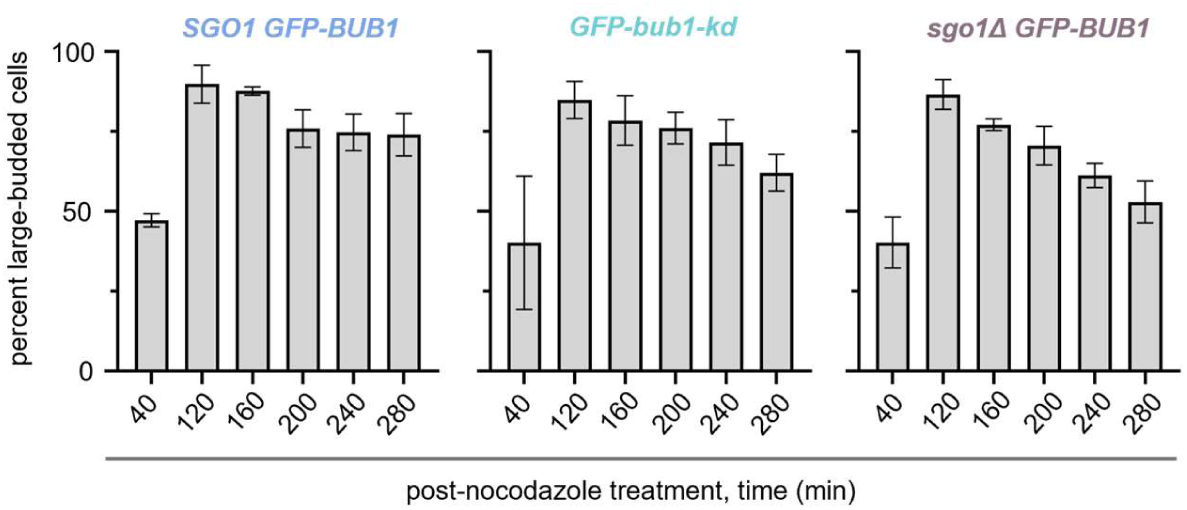
The proportion of large-budded cells scored for the presence or absence of GFP-Bub1 signal post nocodazole treatment. Bar graphs representing the proportion of large-budded cells obtained post nocodazole treatment in the indicated strains, N=3, n>100, cells counted in each experiment. Errors bars represent SD.

**Supplementary Figure 4.**
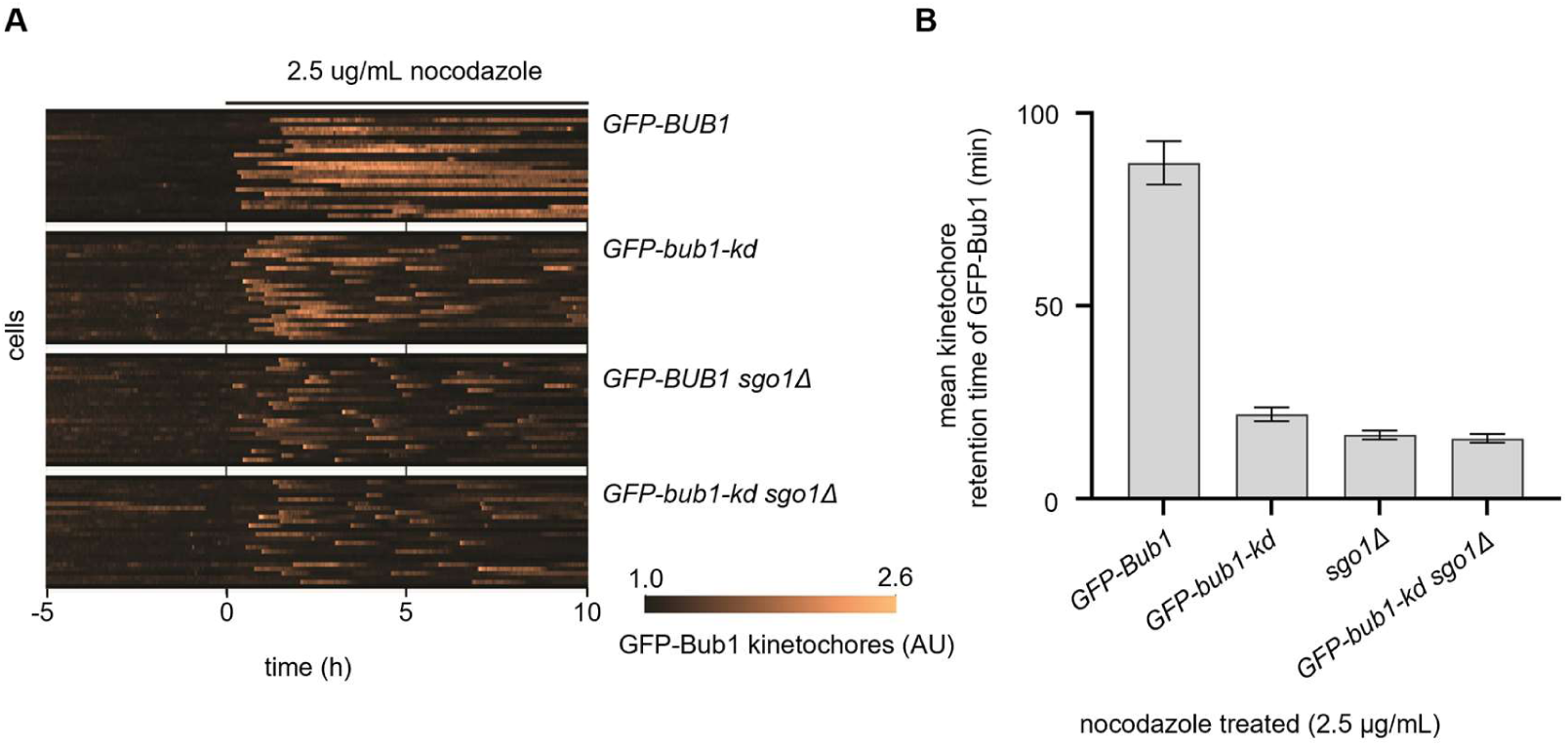
*bub1-kd sgo1Δ* double mutant cells behave similar to single *sgo1Δ* mutant cells. **(A)** Microfluidics assay to determine the retention time of GFP-Bub1 at kinetochores in response to nocodazole treatment. Temporal heat maps of 30 randomly selected cells are shown. The heat map represents the changes in the kinetochore localization signal of GFP-Bub1 over time. Each bright track on the *y*-axis of the heat map represents GFP-Bub1 signals from an individual cell (median of the brightest 5 pixels in each cell divided by the overall cell median brightness). The length of each bright track along the *x*-axis represents the time (min). The time of addition of nocodazole (2.5 μg/mL) is considered as 0 hr. Images are taken every 2 min for 10 h. Assay was performed using IL102 (*GFP-BUB1*), IL143 (*GFP-bub1-kd*), CNSD176 (*GFP-BUB1 sgo1Δ*), and CNSD173 (*GFP-bub1-kd sgo1Δ*) strains. **(B)** Bar graphs representing the quantitative analysis of GFP-Bub1 retention time at kinetochores obtained from microfluidics assays in the above-indicated strains. n=24, error bars represent SEM.

**Supplementary Figure 5.**
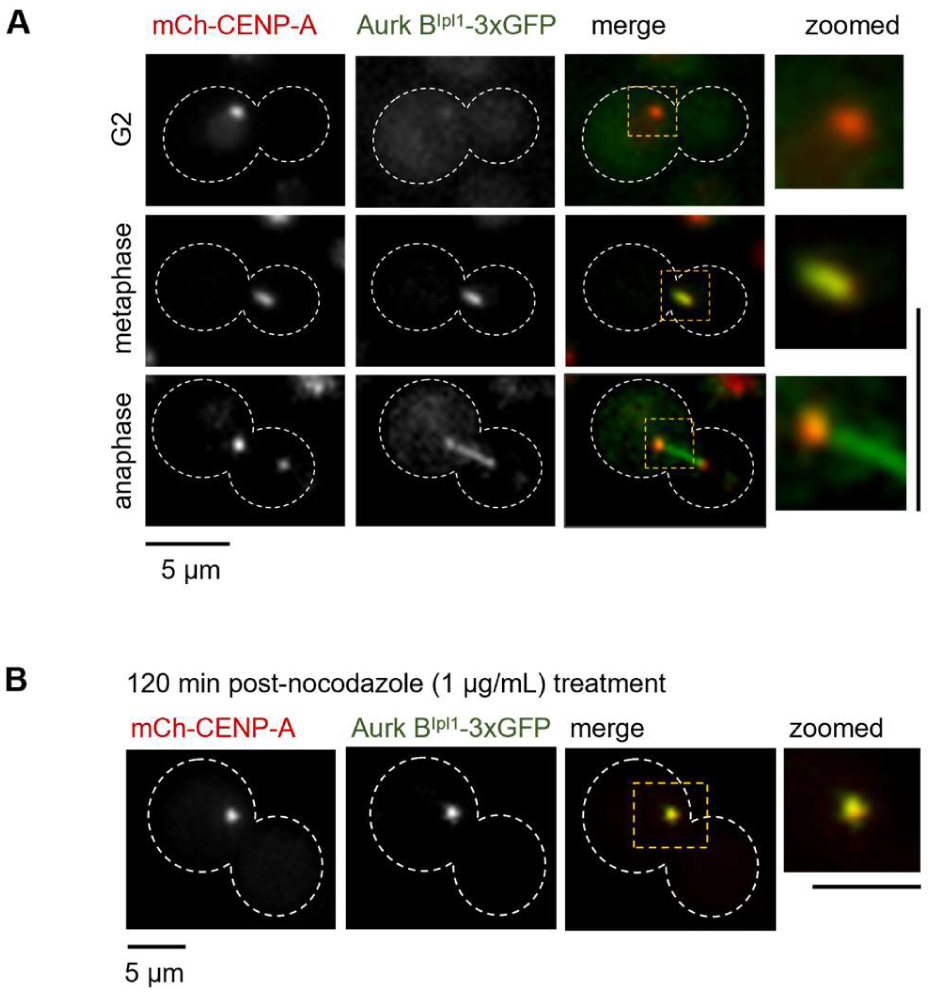
Aurora BIpl1 localizes to the kinetochore (CENP-A) during G2 and mitosis and exhibits spindle-like localization. **(A)** Microscopic images of Aurora BIpl1-3xGFP localized with a kinetochore protein mCherry-CENP-A at G2, metaphase, and anaphase stages of the cell cycle. **(B)** Colocalization of Aurora BIpl1-3xGFP with mCherry-CENP-A when treated with nocodazole (1 μg/mL) for 120 min. Scale bar, 5 μm.

**Supplementary Figure 6.**
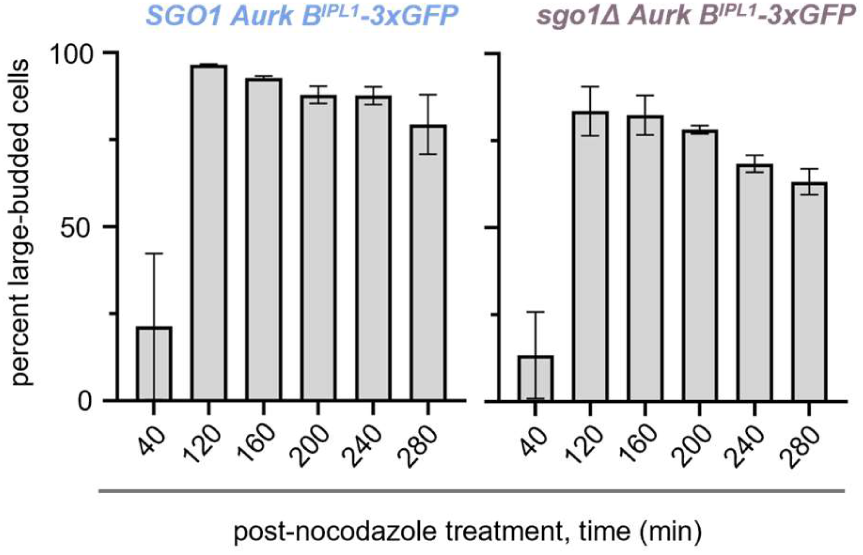
The proportion of large-budded cells scored for the presence or absence of Aurora BIpl1-3xGFP signal post nocodazole treatment. Bar graphs representing the proportion of large-budded cells obtained post nocodazole treatment in the indicated strains, N=3, n>100, cells counted in each experiment. Errors bars represent SD.

**Supplementary Figure 7.**
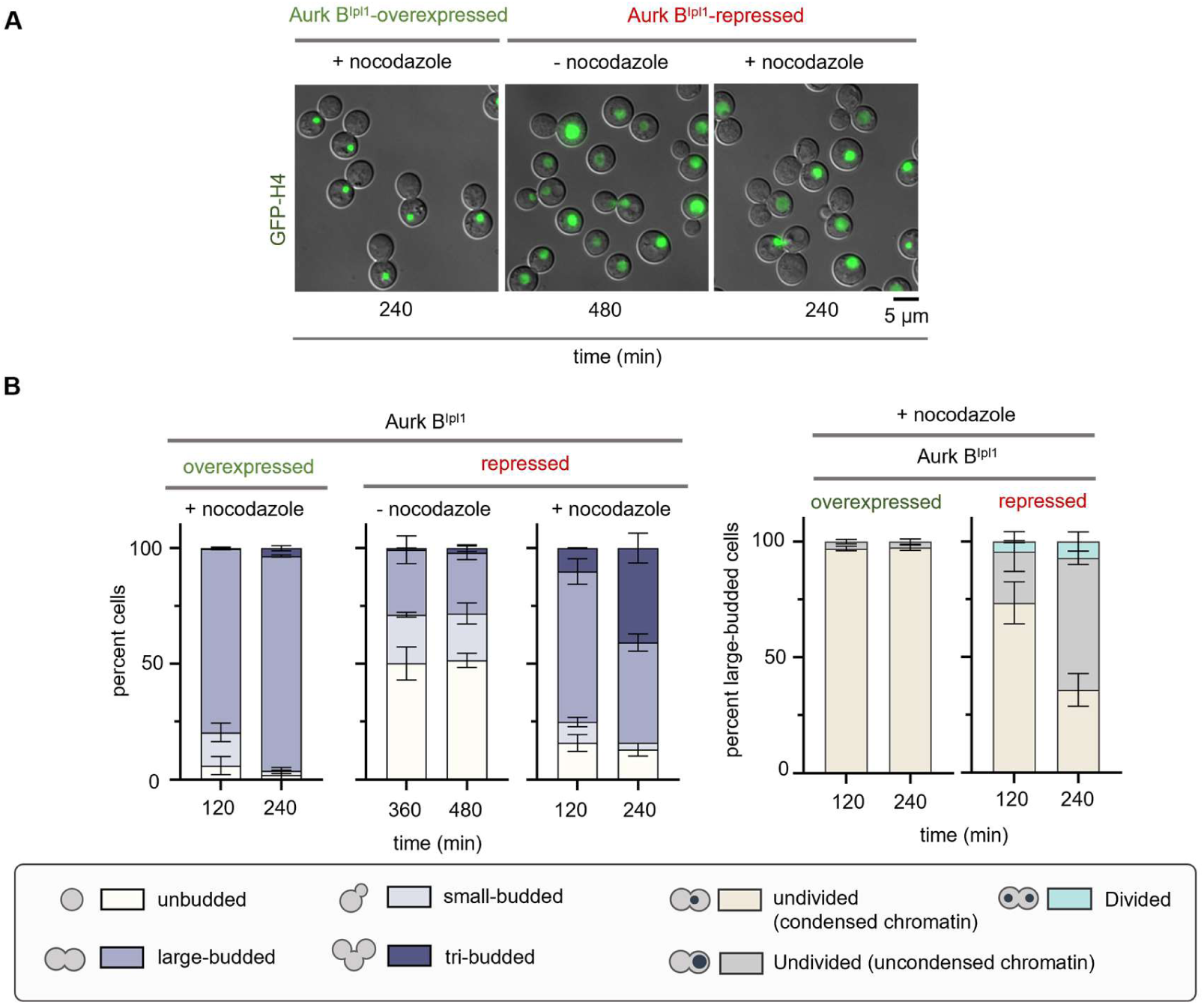
Aurora BIpl1 mutants fail to arrest at metaphase in response to nocodazole treatment. **(A)** Microscopic images of CNNV104 (*GFP-H4 GAL7-AURORA BIPL1*) grown in permissive and non-permissive media in the presence and absence of nocodazole. Scale bar, 5 μm. Representative images of cells treated with nocodazole and depleted of Aurora BIpl1 for 240 and 480 min were shown. **(B)** *Left*, bar graphs representing the proportion of unbudded, small-budded, large-budded, and tri-budded cells. *Right*, bar graphs representing the proportion of large-budded cells of indicated phenotype, N=2, n>100 cells counted for each experiment.

**Supplementary Figure 8.**
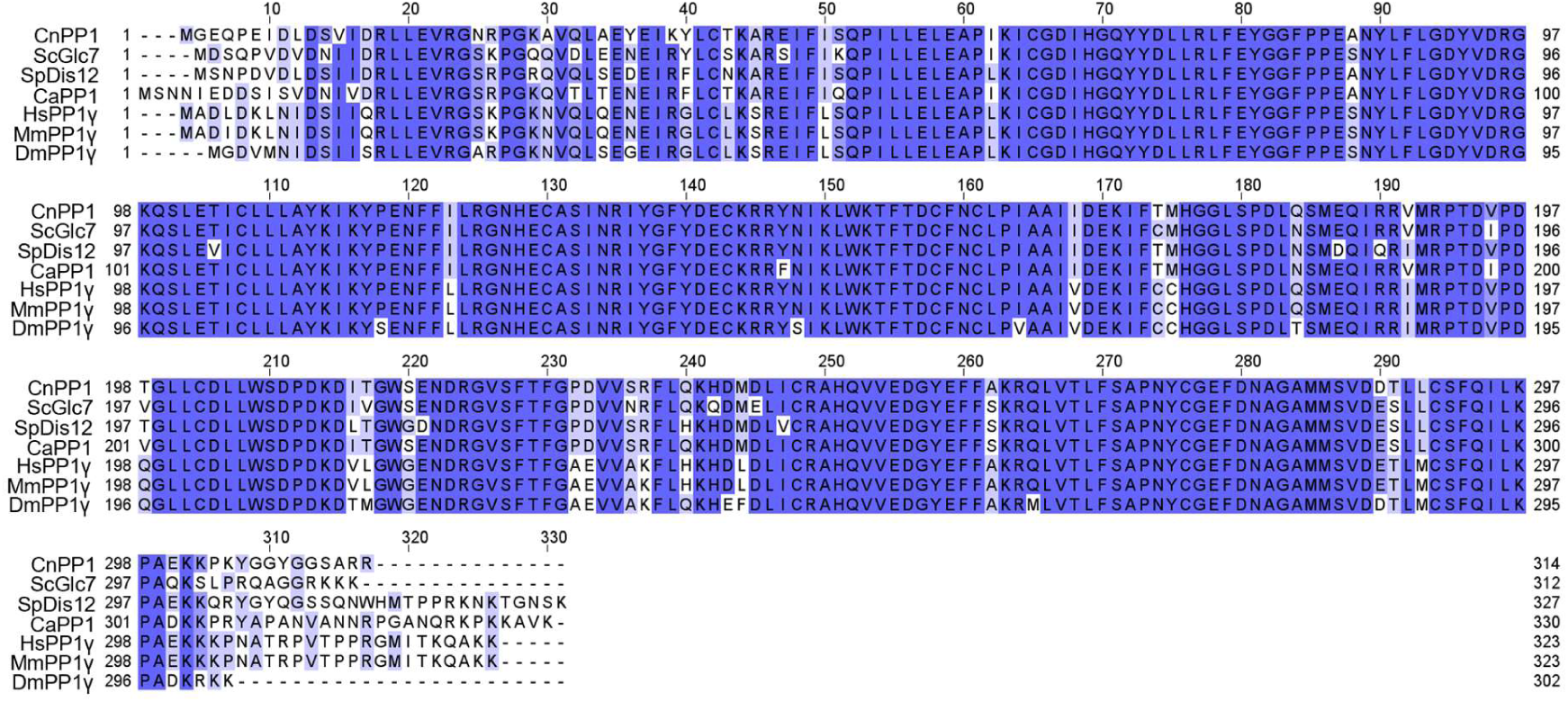
Sequence conservation of *C. neoformans* PP1. Multiple sequence alignment of PP1 homolog of *C. neoformans* (CNAG_03706) with vertebrate and invertebrate species (Cn- *C. neoformans*; Sc- *S. cerevisiae*; Sp- *S. pombe*; Ca- *Candida albicans*; Hs- *H. sapiens*; Mm- *M. musculus*; Dm- *D. melanogaster*) was performed using Clustal Omega (Sievers et al., 2011) and the alignment was formatted using Jalview 2 (Waterhouse et al., 2009). Highly conserved regions are shaded in dark blue.

**Supplementary Figure 9.**
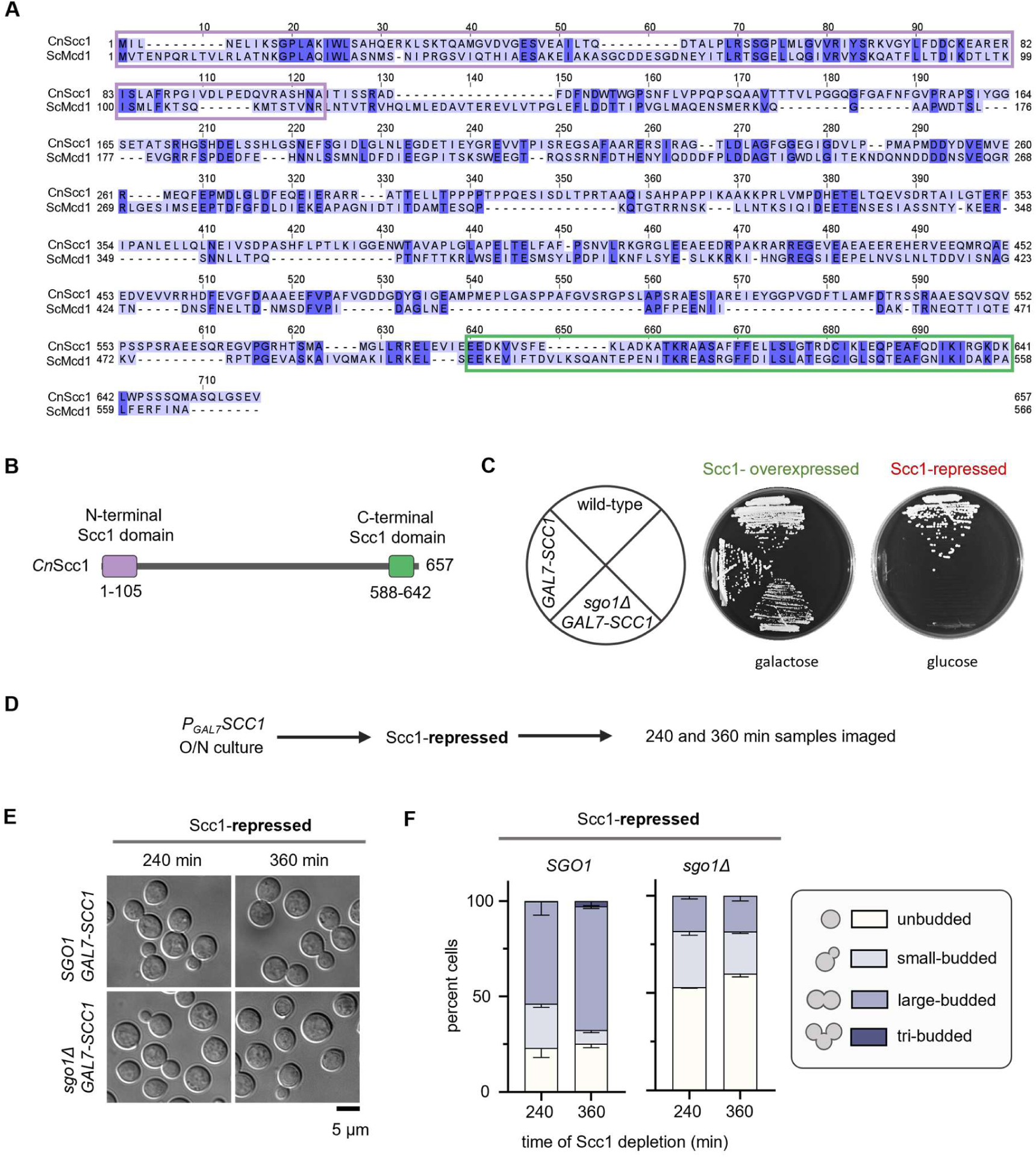
Shugoshin responds to tensionless kinetochore-MT attachments. **(A)** Pairwise sequence alignment of Scc1 homologs of *C. neoformans* (CNAG_01023) and *S. cerevisiae* by Clustal Omega (Sievers et al., 2011), and formatted using Jalview 2 (Waterhouse et al., 2009). Highly conserved regions are shaded in dark blue. The two domains corresponding to N- and C-terminal Scc1 domains (E-values of 2.0e-31 and 2.2e-15, respectively) of CnScc1 are highlighted in violet and green boxes. **(B)** Schematic of the domains present in *C. neoformans* Scc1 protein. **(C)** Plate photographs of strains expressing Scc1 under the regulatable *GAL7* promoter grown in media containing glucose (non-permissive) and galactose (permissive). **(D)** Schematic to check the response of Scc1 repression in *SGO1* and *sgo1Δ* backgrounds. **(E)** Microscopic images of GFP-H4 tagged CNSD181 (*GAL7-3xFLAG-SCC1*) and CNSD182 (*sgo1Δ GAL7-3xFLAG-SCC1*) strains depleted of Scc1. Representative images of cells depleted of Scc1 for 4 and 6 h were shown. Scale bar, 5 μm. **(F)** Bar graphs representing the percentage of unbudded, small-budded, large-budded, and tri-budded cells scored. N=2, n>100 cells for each experiment, error bars represent SD.

**Supplementary Figure 10.**
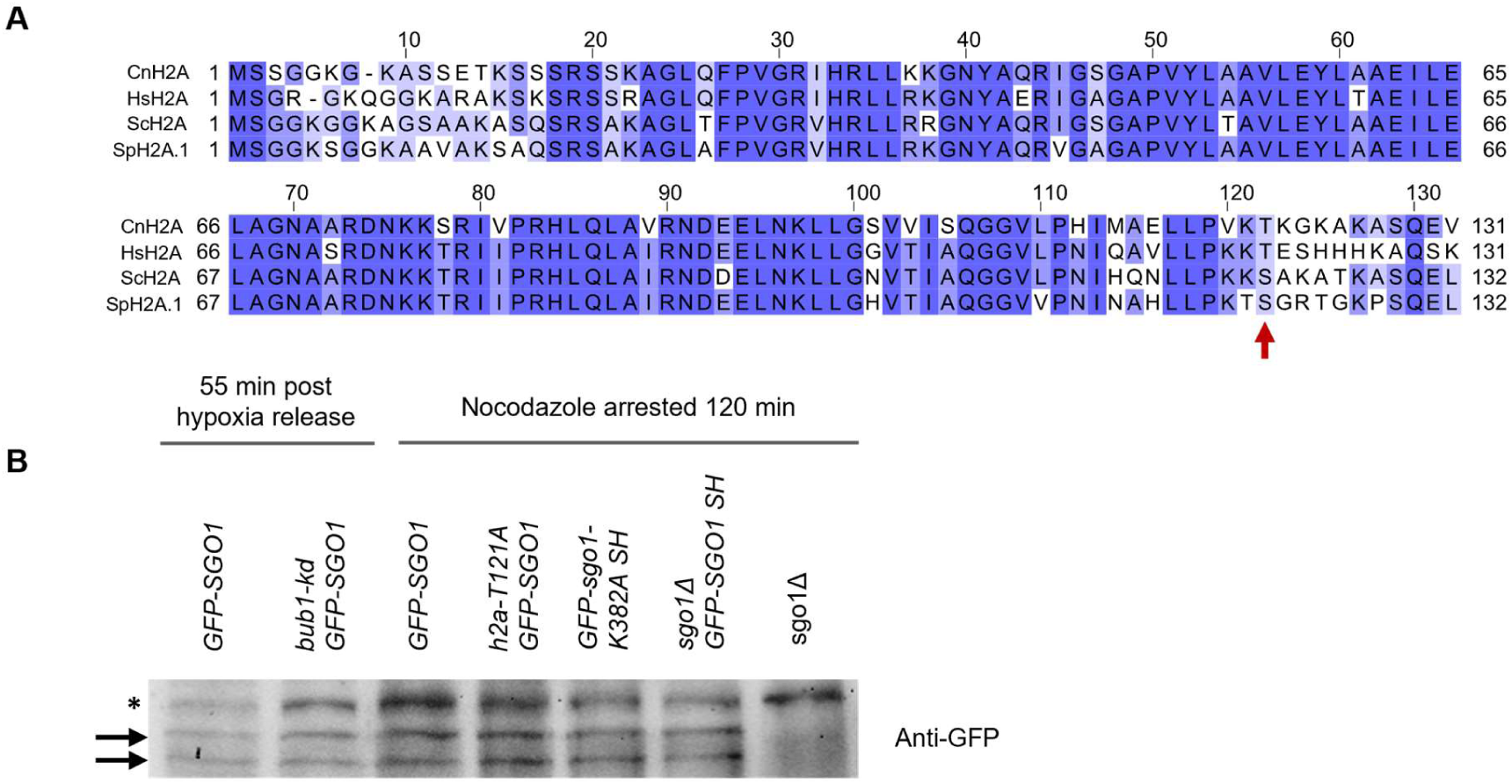
Conservation of histone H2A sequence in *C. neoformans* and expression levels of GFP-Sgo1 in various mutant strains. **(A)** Multiple sequence alignment of H2A homologs obtained from *C. neoformans* (Cn)*, S. cerevisiae* (Sc), *S. pombe* (Sp) and *H. sapiens* (Hs). The alignment was performed using Clustal Omega (Sievers et al., 2011) and formatted using Jalview 2 (Waterhouse et al., 2009). The dark blue shaded regions represent highly conserved amino acid residues. Arrow indicates the conserved T121/S120 residue phosphorylated by Bub1. **(B)** Western blot showing the expression levels of GFP-Sgo1 at metaphase in the indicated strains. * Indicates non-specific band used as a loading control. Arrows indicate two bands of GFP-Sgo1. *sgo1Δ* lane represents no tag control.

**Supplementary Figure 11.**
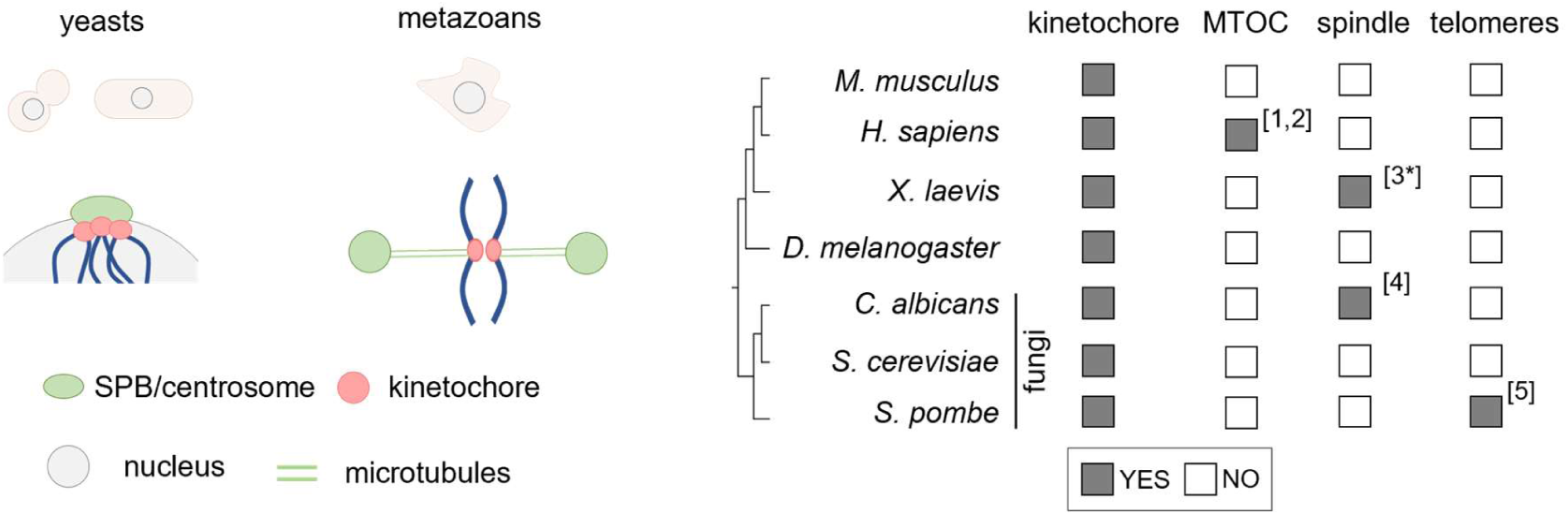
Schematic highlighting the sub-cellular localization of shugoshin reported in various species. *Left*, schematic highlighting the spatial positioning of SPB/centrosome and centromeres in yeasts and metazoans. *Right*, localization of shugoshin with respect to the kinetochore, MTOCs (centrosomes or SPBs), spindle MTs, and telomeres. Filled and open squares represent the presence or absence of shugoshin at the indicated sub-cellular locations. The data was compiled from reviews (Marston, 2015, Gutierrez-Caballero et al., 2012). [1,2] (Mohr et al., 2015, Wang et al., 2008), [3*] (Salic et al., 2004), this study has shown that the purified N-terminal region of shugoshin is capable of binding to MTs *in* vitro. [4] (Sane et al., 2021) and [5] (Tashiro et al., 2016).

**Table S1:**
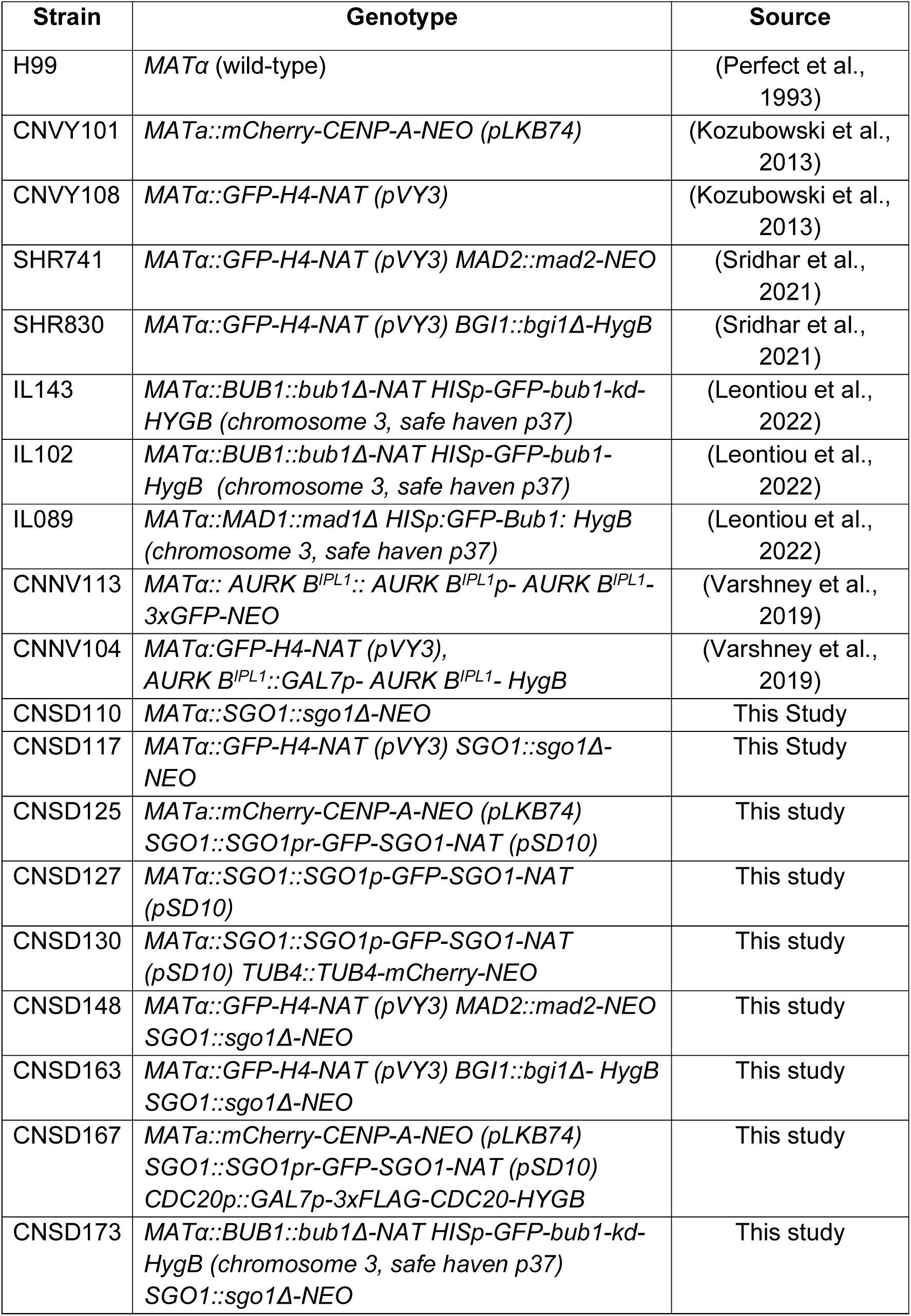
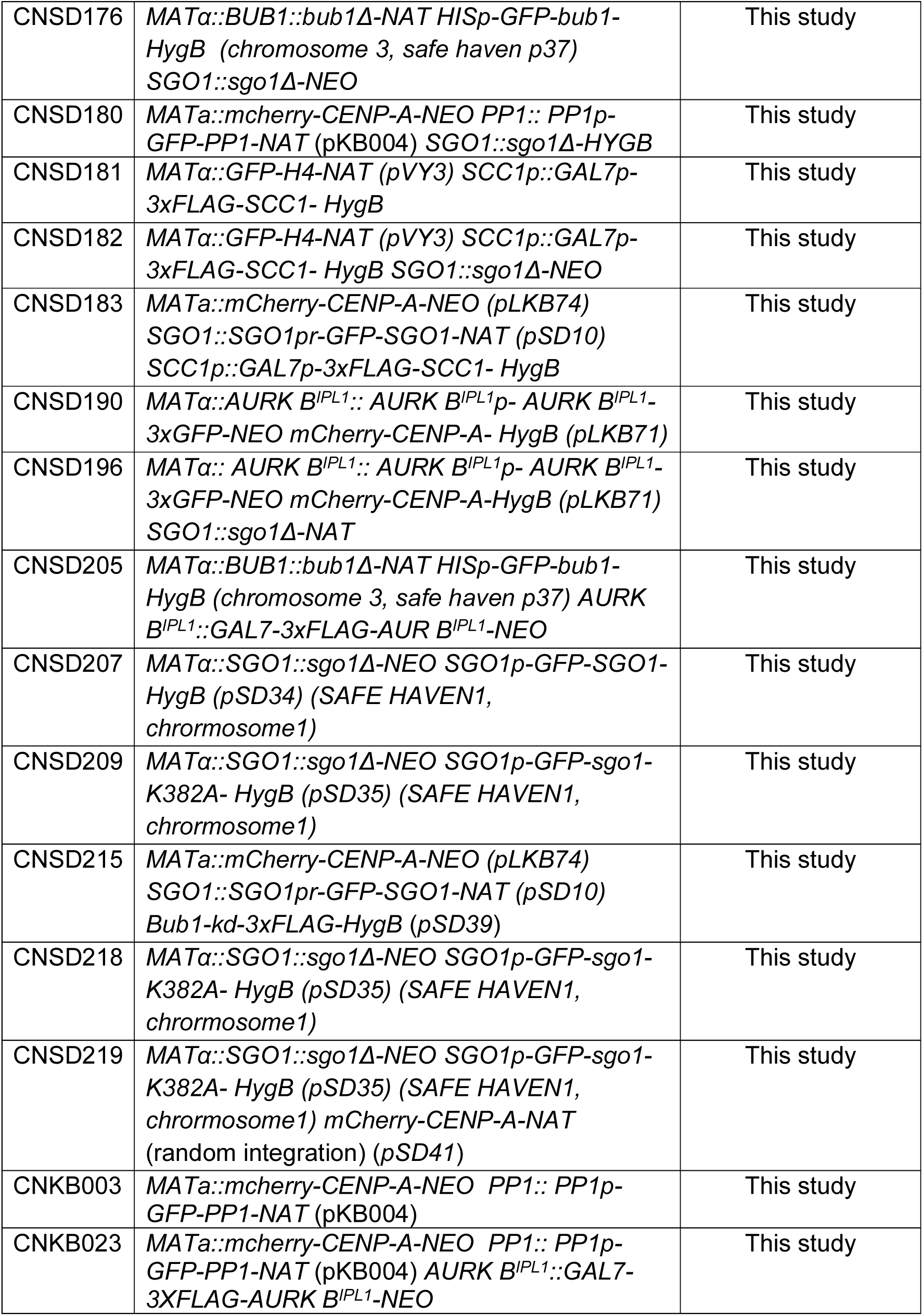
List of strains.

**Table S2:**
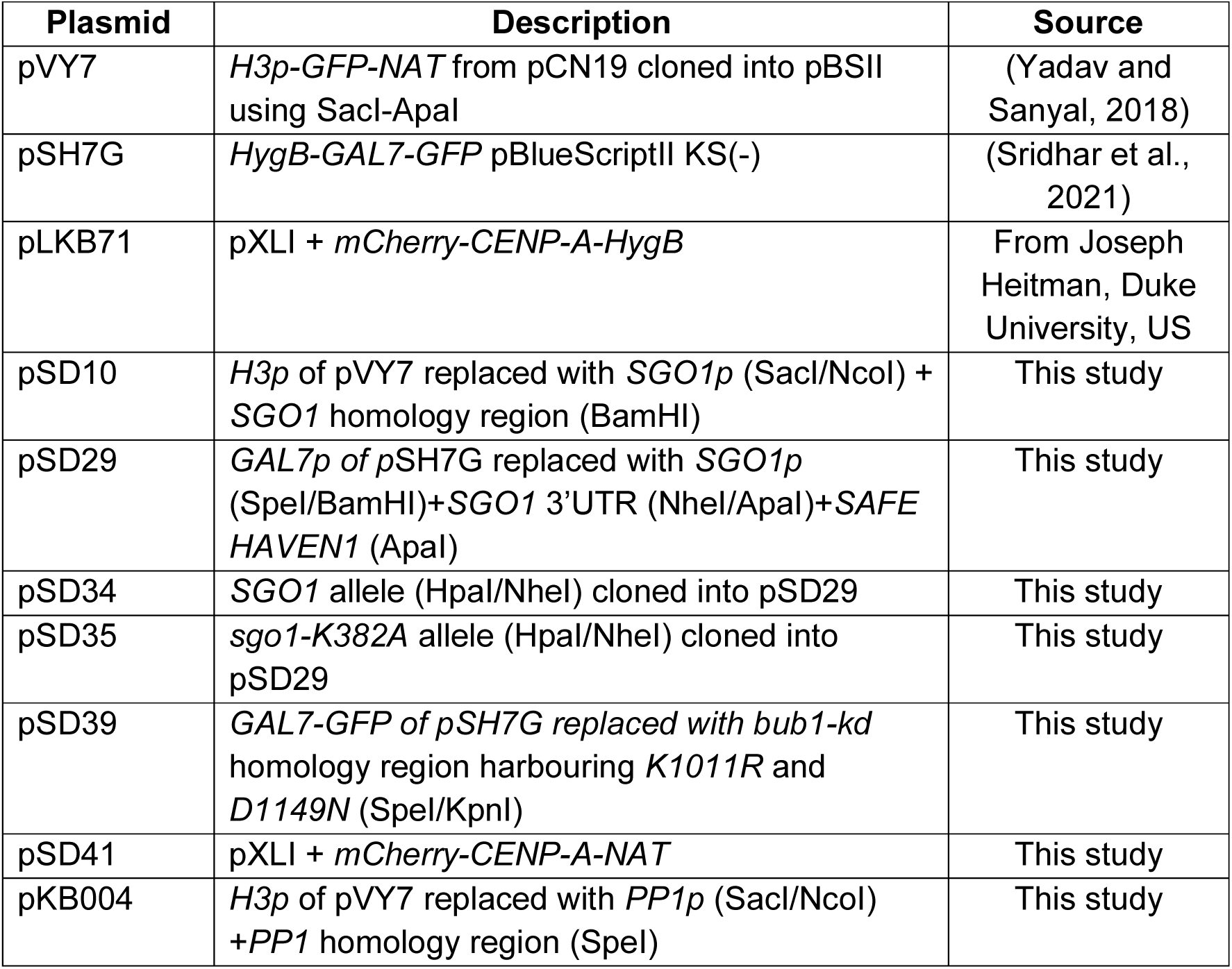
List of plasmids.

| Plasmid | Description | Source |
| --- | --- | --- |
| pVY7 | *H3p-GFP-NAT* from pCN19 cloned into pBSII using *SacI*-*Apal* | (Yadav and Sanyal, 2018) |
| pSH7G | *HygB-GAL7-GFP* pBlueScriptII KS(-) | (Sridhar et al., 2021) |
| pLKB71 | pXLI + *mCherry-CENP-A-HygB* | From Joseph Heitman, Duke University, US |
| pSD10 | *H3p* of pVY7 replaced with *SGO1p* (*SacI*/*NcoI*) + *SGO1* homology region (*Bam*HI) | This study |
| pSD29 | *GAL7p* of pSH7G replaced with *SGO1p* (*SpeI*/*Bam*HI)+*SGO1* 3'UTR (*NheI*/*Apal*)+*SAFE HAVEN1* (*Apal*) | This study |
| pSD34 | *SGO1* allele (*HpaI*/*NheI*) cloned into pSD29 | This study |
| pSD35 | *sgo1-K382A* allele (*HpaI*/*NheI*) cloned into pSD29 | This study |
| pSD39 | *GAL7-GFP* of pSH7G replaced with *bub1-kd* homology region harbouring *K1011R* and *D1149N* (*SpeI*/*KpnI*) | This study |
| pSD41 | pXLI + *mCherry-CENP-A-NAT* | This study |
| pKB004 | *H3p* of pVY7 replaced with *PP1p* (*SacI*/*NcoI*) +*PP1* homology region (*SpeI*) | This study |

**Table S3:**
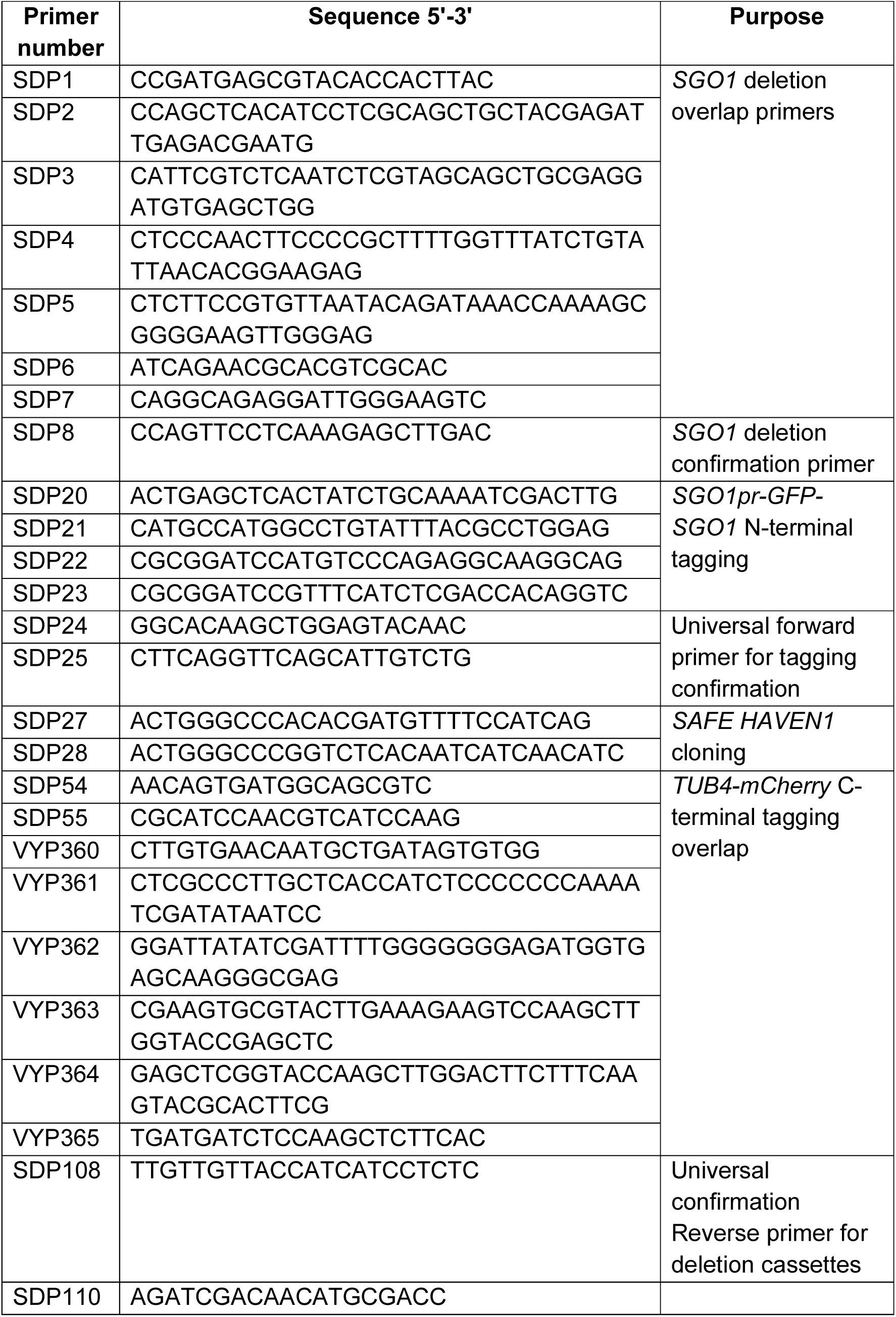
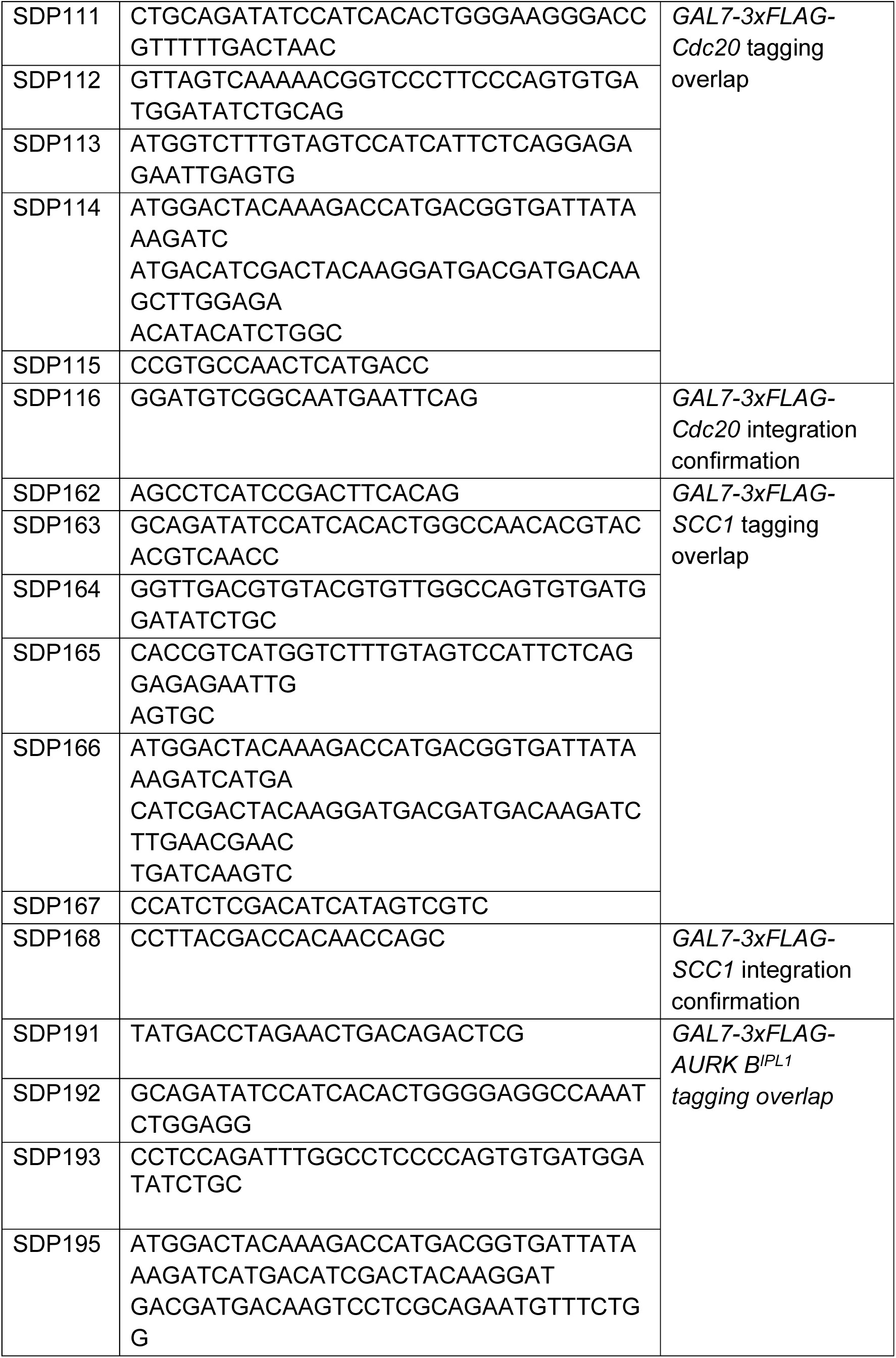
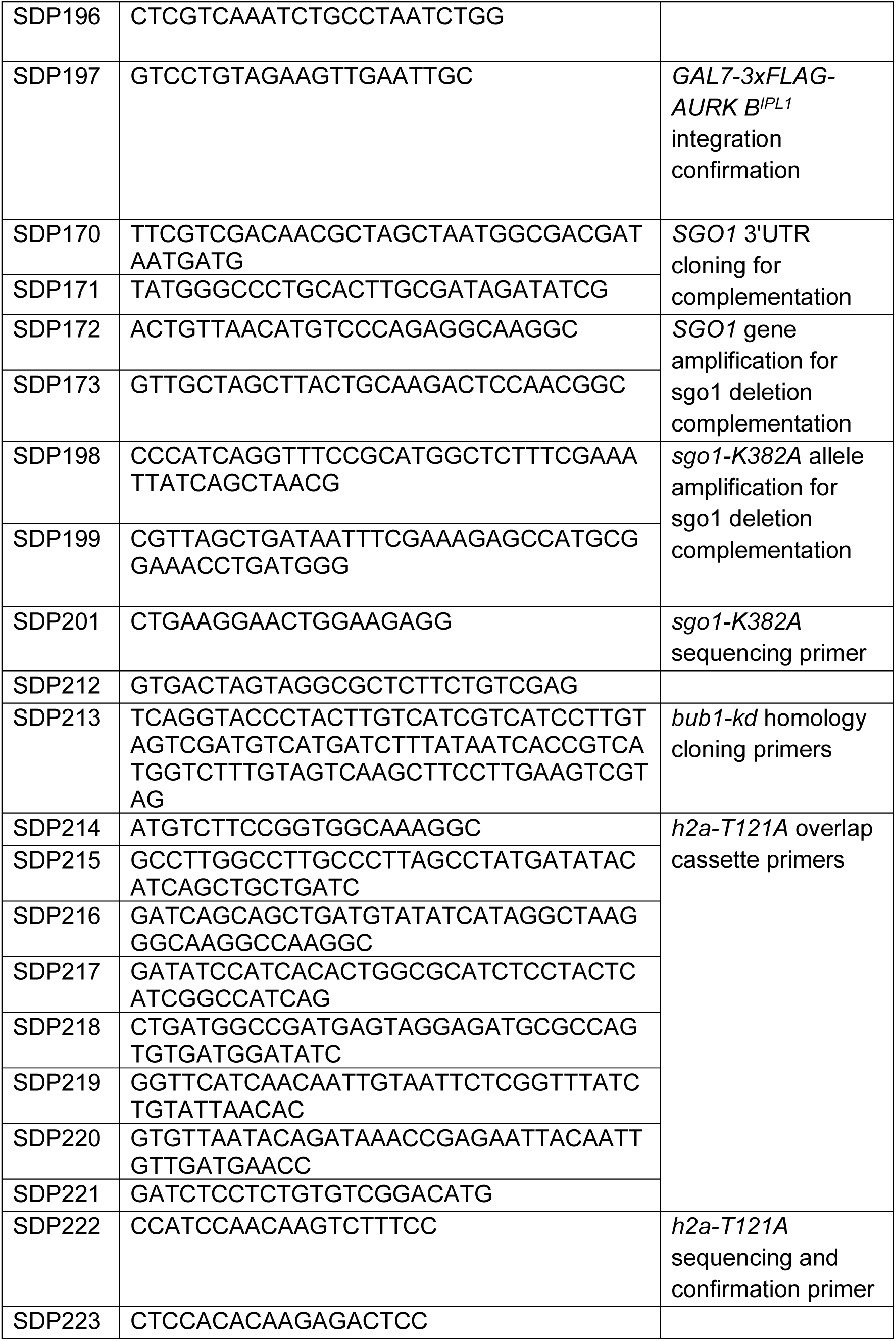
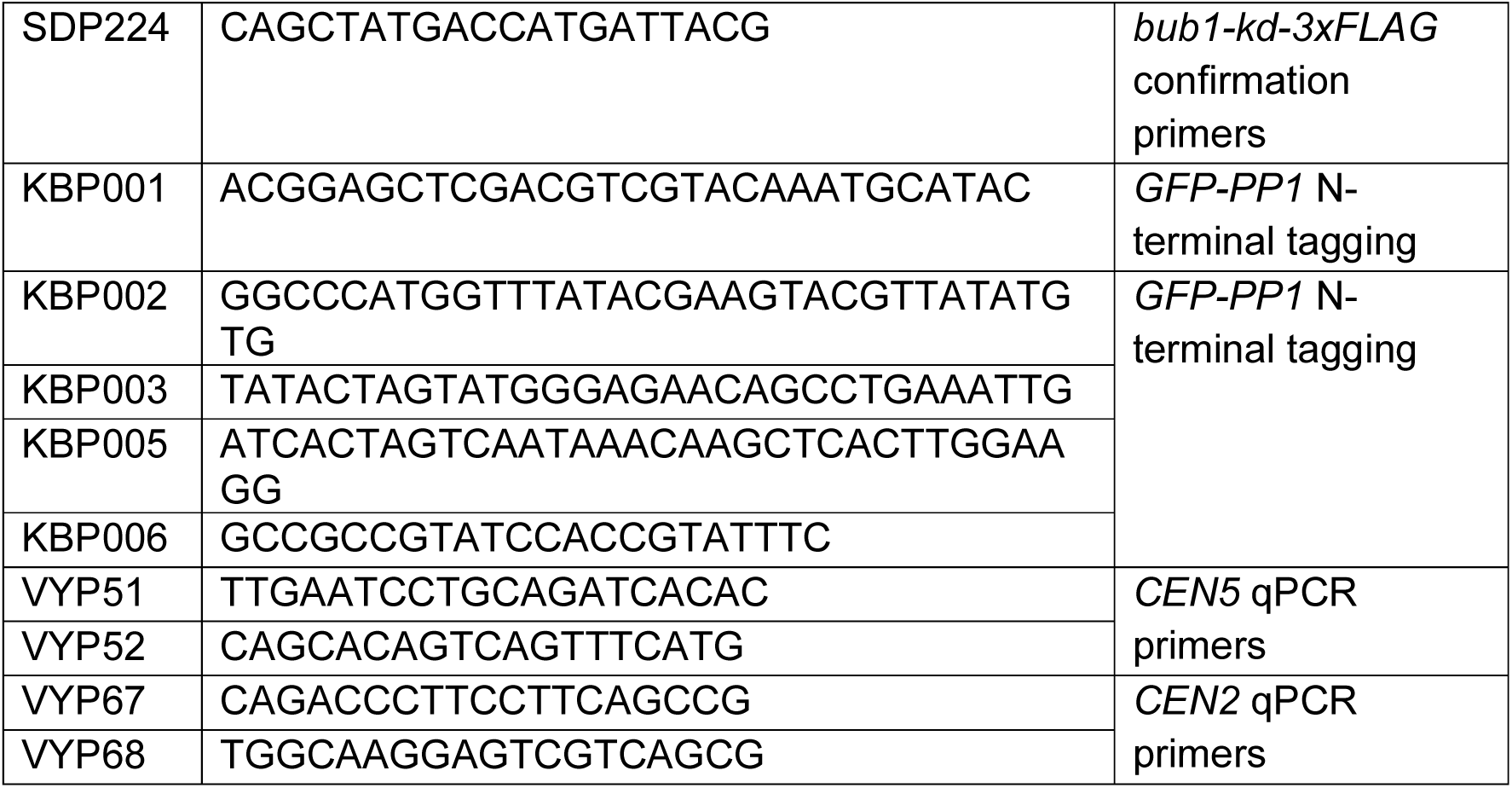
List of primers.

